# DDX3X syndrome mutations lock DDX3X-RNA conformational states to drive persistent pathological condensation and neuronal death

**DOI:** 10.64898/2026.01.29.702492

**Authors:** Poulami Ghosh, Shivani Krishna Kapuganti, Sabhyata Gopal, Swati Lamba, Purusharth I Rajyaguru, Sannula Kesavardhana

## Abstract

DDX3X is a highly conserved RNA helicase associated with RNA metabolism, translation initiation, and deciding cell fate choices. Spontaneous mutations in DDX3X cause a rare genetic human disorder called DDX3X syndrome, showing a spectrum of neurodevelopmental and intellectual abnormalities. How missense mutations in DDX3X lead to aberrant cellular functions and pathological consequences is unclear. Here, we demonstrate that specific DDX3X syndrome missense mutations induce the formation of persistent, solid-like DDX3X stress granules by structurally altering the conformation of the DDX3X-RNA complex. Structural interrogation of DDX3X syndrome missense mutations revealed critical mutations that perturb the DDX3X-RNA complex, exhibit high clinical pathogenicity scores, and are associated with cancers. N and C-terminal mutations adjacent to the helicase domain of the DDX3X (F182V, I190S, T198P, L556S, and L559H) showed augmented stress granule (SG) assembly, and formation of persistent DDX3X-SGs in neuronal and non-neuronal cells. Intriguingly, these mutations altered the liquid-like properties of DDX3X-SGs, forming solid-like SGs. Mechanistically, these mutations drive the persistent DDX3X-SGs by either locking DDX3X in an open, RNA-bound conformation or by increasing the rigidity of the DDX3X-RNA complex (I190S; T198P), thereby conferring a loss of liquid-like properties. The persistent solid-like DDX3X-SGs preferentially promoted neuronal lytic cell death and did not affect translation. Also, the C-terminal L556S and L559H mutations, which form persistent granules, promoted the DDX3X-driven β-amyloid aggregation. Our observations indicate that DDX3X syndrome mutations near the N- and C-termini promote the formation of solid-like DDX3X condensates, neuropathological aggregation, and neuronal cell death, which might underlie the disease pathogenesis in humans.

**GRAPHICAL ABSTRACT:** 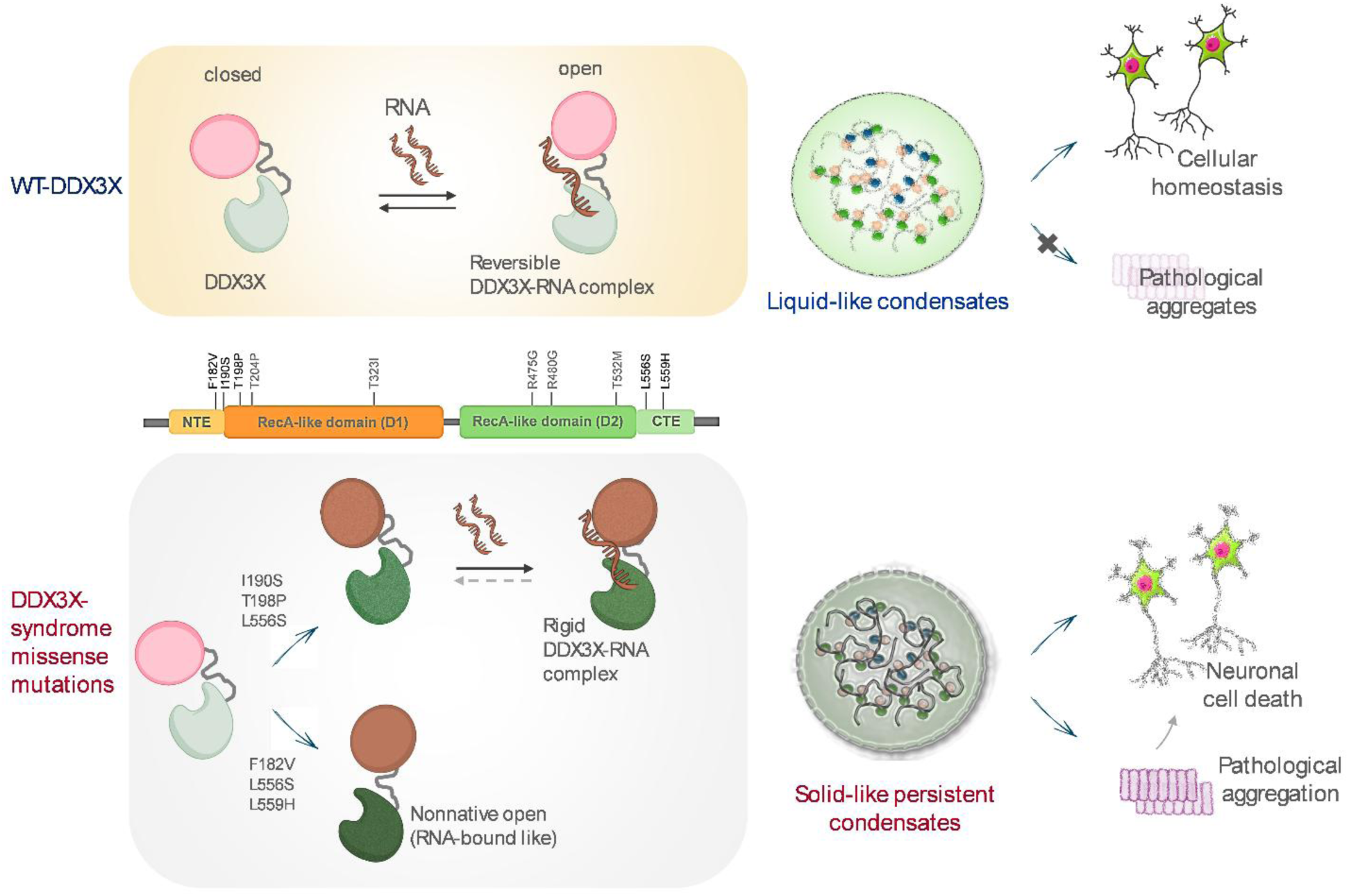

## INTRODUCTION

DDX3X is an X-linked DEAD box protein with RNA helicase function, regulating diverse cellular functions, including mRNA metabolism, translation initiation, innate immune activation, and critical cell fate choices determining development and tumorigenesis ^1–9^. In response to diverse cellular stress conditions, DDX3X promotes the assembly of stress granules (SGs) to stall the translation transiently and enable cell survival, and upon removal of stress, the SGs disassemble ^2; 4; 9; 10^. In addition to translation and SGs regulation, DDX3X orchestrates innate immune activation and cell fate choices ^4; 7; 8; 11^. During stress conditions, DDX3X promotes the assembly of the NLRP3 inflammasome, which in turn activates cell fate towards pyroptosis, an inflammatory form of cell death. Additionally, DDX3X regulates immune cell fate decisions by modulating the crosstalk between SGs and the NLRP3 inflammasome ^4; 8^.

DDX3X consists of a core helicase domain with two RecA-like domains (D1 and D2) and the flanking N and C-terminal intrinsically disordered regions (IDRs) ^11; 12^. The core helicase of DDX3X is required for ATP binding and its subsequent hydrolysis, as well as for binding to RNA, which facilitates the resolution of complex RNA structures. The N-and C-terminal IDRs of DDX3X promote liquid-liquid phase separation (LLPS) for assembling SGs ^5; 11; 13; 14^. Also, the C-terminal extension near the helicase domain is known to promote oligomerization of DDX3X ^11; 12^. The conformationally dynamic nature of the DDX3X in RNA-bound is critical for its function in LLPS and to restrict the solid-like phase condensates, which might be associated with loss of cellular functions and disease pathogenesis.

During early developmental stages, DDX3X is expressed highly in cortical progenitor cells and neurons and is vital for healthy hindbrain development, corticogenesis, and synaptogenesis ^15–17^. DDX3X syndrome is a sporadic, rare genetic neurodevelopmental disorder due to mutations in the DDX3X protein, leading to intellectual disabilities, neuronal, sensory, movement, and behavioral abnormalities ^18–21^. DDX3X syndrome is predominantly seen in females, where mutations in DDX3X cause developmental delays and intellectual disability, but some of the patients survive to the adult stage ^16; 21; 22^. DDX3X syndrome is rare in males and is detrimental to the survival of males because of the presence of a single X chromosome, and can lead to profound neurodevelopmental impairment ^18; 19^. In addition, mutations in human DDX3X are associated with enhanced susceptibility to certain cancers ^15; 16; 23^. The clinical pathologies and the anatomical and developmental abnormalities of the DDX3X syndrome are well studied ^16; 17; 21; 22^, however, how mutations in DDX3X alter cellular functions and cause a spectrum of neuronal abnormalities are unclear. Particularly, whether the missense mutations in DDX3X syndrome patients confer loss or gain of function to disrupt the DDX3X-RNA complex functions regulating SGs and neuronal cell fate choices have not been studied. Pathological aggregates, like amyloid-β and Tau depositions, disrupt neuronal and microglial physiology and cause neurodegeneration and neuroinflammation ^24–27^. It remains to be established whether mutations in DDX3X promote neuropathological aggregation and neurodevelopmental abnormalities.

In this study, we defined specific DDX3X syndrome mutations that impact the structures of DDX3X-RNA complexes and show high pathological scores. Structural, functional, and mechanistic studies show that specific DDX3X syndrome mutations spanning the N and C-terminal regions of the DDX3X promote persistent solid-like DDX3X condensates, driving neuronal cell death and amyloid aggregation. Our study provides specific mechanisms by which these DDX3X syndrome mutations alter the dynamics of DDX3X, with and without RNA binding, to drive condensate formation, loss of liquid-like properties, and the formation of pathological, persistent granules in cells.

## RESULTS

### Structure-based evaluation identified highly pathogenic DDX3X missense mutations

Since the first report was published in 2015 ^21^, several studies have identified DDX3X mutations associated with the DDX3X syndrome. Various missense, nonsense, and frameshift mutations have been identified in patients with DDX3X syndrome **(Figure 1A)**. Among the mutations reported so far, we sought to identify missense mutations that would impact the three-dimensional structure and RNA-binding interface of DDX3X, potentially altering DDX3X function. Analysis of the residue accessibility, inter-residue association, and RNA binding interface residues in the 3D structure of the DDX3X-RNA complex revealed a few DDX3X syndrome mutations (F182V, I190S, T198P, T204P, T323I, R475G, R480G, T532M, L556S, and L559H) which span helicase domain and the disordered N-terminal extension (NTE) and C-terminal extension (CTE) regions, that likely alter DDX3X structure and function **(Figure 1B-C**). AlphaMissense-based clinical pathogenicity prediction analysis ^28^ revealed that the selected DDX3X syndrome mutations appeared to be clinically highly pathogenic, further corroborating the significant impact of the selected mutations on protein structure and clinical function **(Figure 1D)**. In addition, these structure-based rationally selected residues of the DDX3X syndrome were mutated in various cancers, suggesting the critical role of these positions in DDX3X function **(Figure 1E)**.

**Figure 1.**
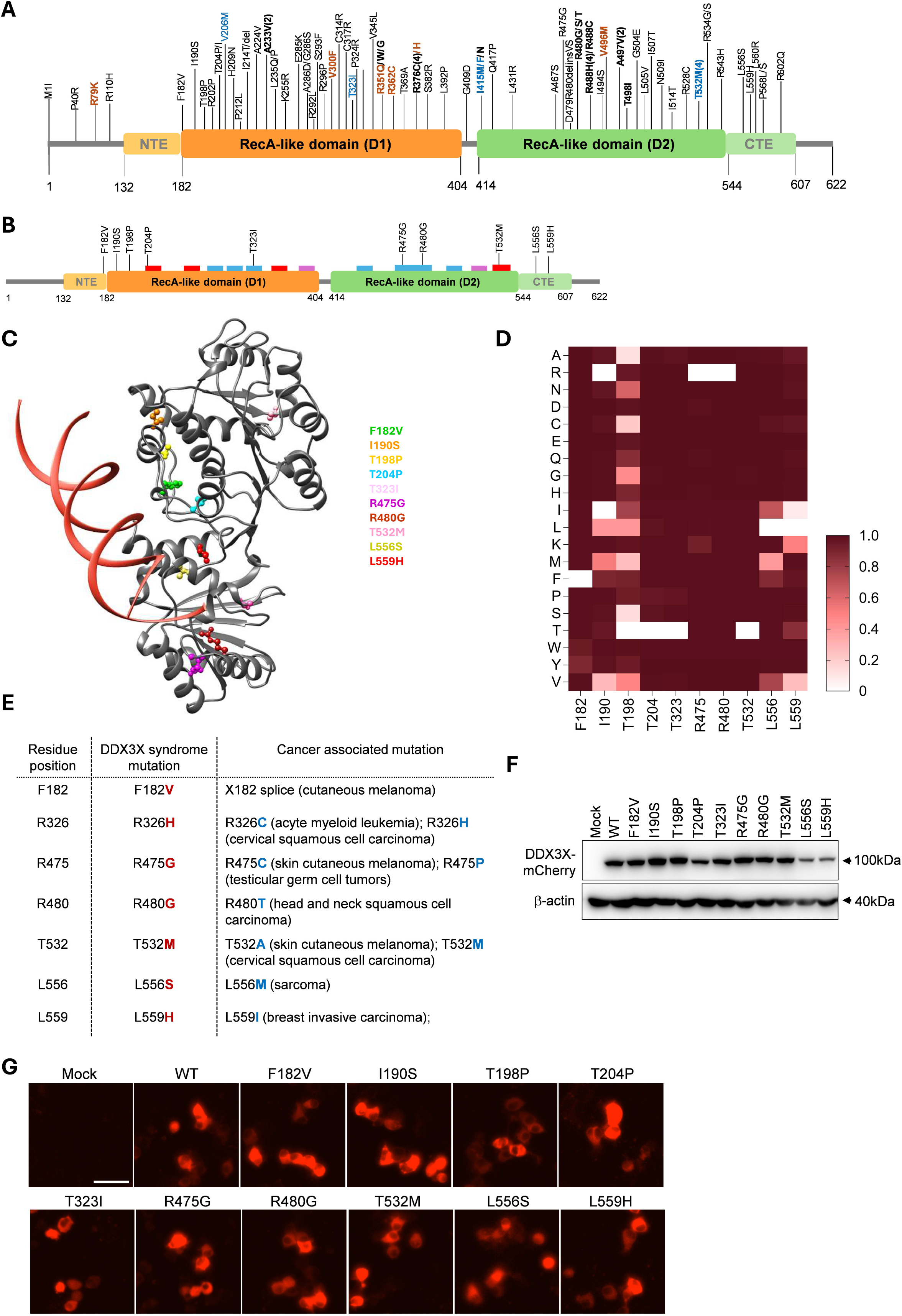
Structure-guided evaluation of DDX3X missense variants to establish pathogenic variants. **A**. Schematic representation of DDX3X protein showing pathogenic mutations identified in DDX3X syndrome patients (bold- multiple occurrences, blue- polymicrogyria, brown- male patients). **B**. Schematic representation of DDX3X functional domains indicating critical DDX3X syndrome-associated mutations selected based on their predicted structural impact. Blue, RNA binding sites; Red, ATP binding and hydrolysis regions; Magenta, regions of interaction between ATP binding and RNA binding residues. **C**. Ribbon structural representation of DDX3X (dark grey) bound to dsRNA (red) (PDB ID: 6O5F), displaying surface accessibility and RNA interface proximity of DDX3X syndrome mutations. Mutated residues are color-coded in the represented structure. **D**. Heat map depicting the AlphaMissense prediction for the likely pathogenicity of selected DDX3X syndrome missense mutations, with dark red being the likely pathogenic and white being the likely benign mutation. **E**. Table depicting selected mutations in DDX3X that have been associated with different cancer types. **F**. Immunoblotting analysis of DDX3X showing the impact of DDX3X syndrome mutations on protein expression in Neuro2a cells. **G**. Fluorescence images of N2a cells expressing DDX3X-mCherry with DDX3X syndrome mutations, captured using Incucyte imager.

To examine whether the selected DDX3X mutations affect DDX3X expression, we monitored intracellular expression of DDX3X mutations using DDX3X-mCherry and found that DDX3X syndrome mutations did not alter DDX3X expression at the cellular level **(Figure 1G)**. Furthermore, the immunoblotting analysis indicated that DDX3X syndrome mutations did not significantly alter protein expression, except for the C-terminal mutations, L556S and L559H **(Figure 1F).** Thus, the selected DDX3X syndrome mutations might alter DDX3X function without affecting its expression.

### DDX3X syndrome mutations augment SG formation

DDX3X regulates cell fate choices by orchestrating the assembly of biological condensates ^4; 8; 9; 29^. Upon stress stimuli, DDX3X phase separates and forms SGs to facilitate translational arrest and retain a cell survival state, and loss of DDX3X significantly diminishes SG formation ^4; 5^. To study the effects of DDX3X syndrome mutations on SG formation, we attempted to generate HeLa cells lacking DDX3X to reconstitute with the DDX3X mutant expressing constructs **(Figure S1A)**. However, these cells were not sustained in culture, and only cells that had DDX3X expression survived and propagated. The selected mutations were ectopically expressed in WT-HeLa cells and subjected to sodium arsenite (SA) treatment to induce SG formation. All the mutant-expressing cells showed the formation of cytoplasmic SGs, in which DDX3X colocalized with G3BP1, a hallmark of SGs **(Figure S1B-E)**. However, the I190S, T198P, L556S, and L559H mutations significantly augmented the number of DDX3X-SGs as compared to the WT-DDX3X **(Figure S1C).** Also, the R480G and T532M mutations showed increased SG formation compared with WT-DDX3X, although this was not significant. To understand whether the increased SG formation phenotype of DDX3X mutations is cell type specific, we expressed DDX3X mutants in neuro 2a (N2a) cells and monitored SG quantification **(Figure S2A-D).** We found that F182V, I190S, T198P, L556S, and L559H mutations significantly augmented DDX3X-SG formation in N2a cells **(Figure S2B)**. Notably, these mutations, either adjacent to the NTE (F182V, I190S, and T198P) or in the CTE (L556S and L559H) regions of DDX3X, augmented SG formation, unlike the helicase domain mutations. These observations indicate that specific pathogenic DDX3X syndrome mutations increase the number of SGs assembled in neuronal and non-neuronal cells.

### DDX3X syndrome mutations in the vicinity of NTE and CTE regions form persistent solid-like SGs

Aberrant aggregation of cellular liquid-like compartments is associated with disease pathogenesis ^30; 31^. Prolonged persistence of SGs is known to favor conversion of liquid-like SGs into solid-like SGs, which, in turn, potentially alter cell-fate choices ^13; 30–32^. To understand whether DDX3X syndrome mutations favor SG persistence in cells, we established an SG wash-off assay by treating cells with SA, followed by wash-off and imaging for SG persistence (**Figure 2A).** WT-DDX3X expressing HeLa cells showed SG formation after SA treatment, and these granules disassembled after washing off SA. Intriguingly, T198P, T323I, and L556S DDX3X mutants showed persistence of SGs in HeLa cells after washing off SA, indicating that these mutations promote the formation of persistent DDX3X-SGs (**Figure 2B-D).** R480G mutant appeared to promote persistent SGs, preferably G3BP1-specific SGs. To further corroborate this data, we performed a wash-off assay in N2a cells. We observed that SGs in WT DDX3X-expressing cells were dissolved after washing off SA, whereas DDX3X mutants, including I190S, T198P, T323I, L556S, and L559H, promoted the persistence of SGs in N2a cells even after SA wash-off **(Figure 2E-G).** Thus, DDX3X syndrome mutations, in particular NTE and CTE mutations, alter DDX3X function to promote persistent SG formation. The R480G and T323I mutations span the core helicase domain and appeared to have a cell-type-dependent effect on persistent SG formation.

**Figure 2.**
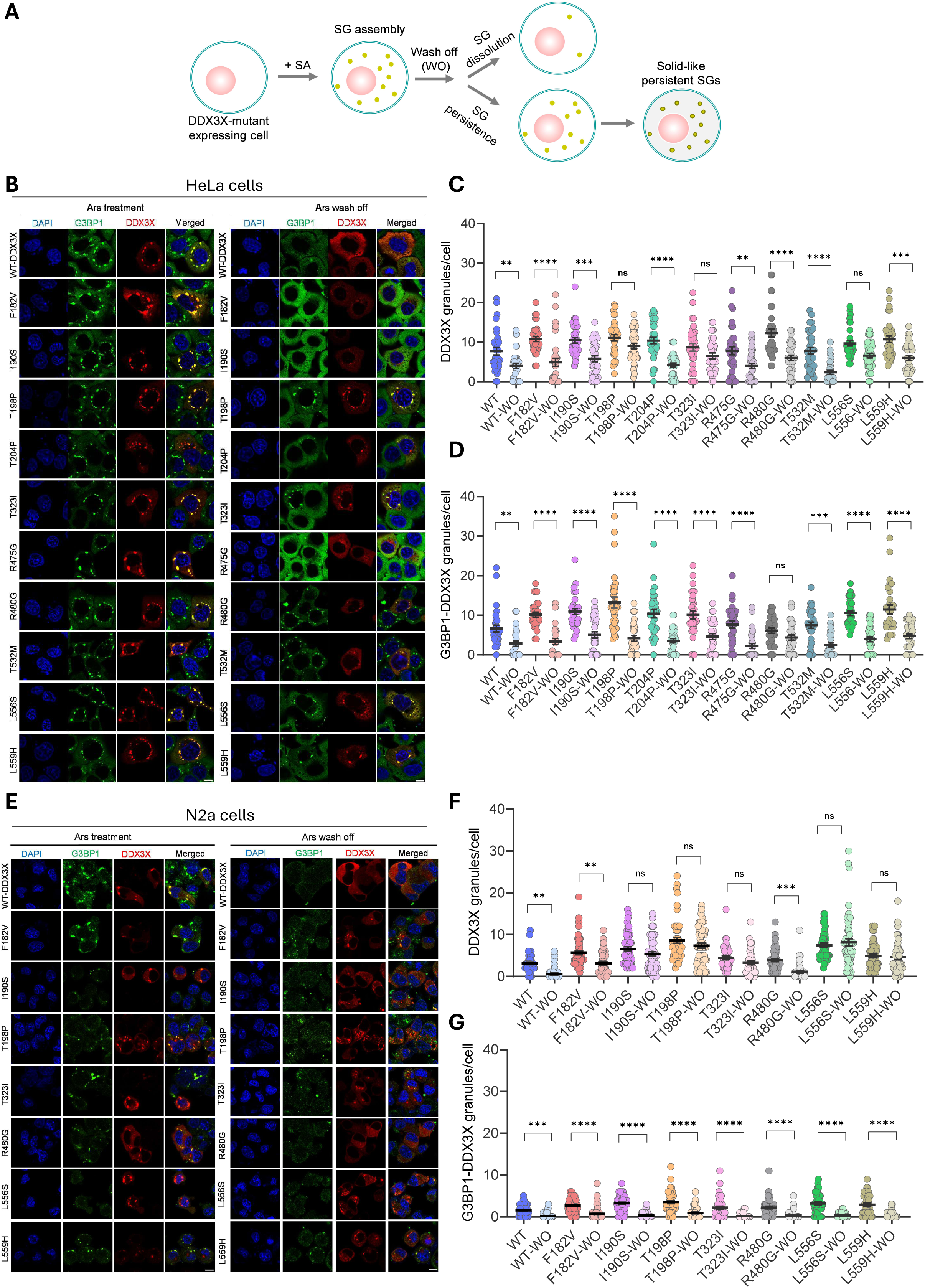
DDX3X missense mutations promote persistent DDX3X-stress granules in neuronal and non-neuronal cells. **A**. Schematic showing the protocol used for monitoring persistent granules in cells. Cells were treated with sodium arsenite (SA), followed by removing SA and supplementing fresh media (wash-off, recovery period) to monitor persistent granules. **B**. Representative confocal microscopy images of HeLa cells expressing WT or mutant DDX3X-mCherry showing stress granules (SGs) upon SA treatment and after wash-off (recovery), immunostained for G3BP1 (green) and DAPI (blue). Scale bar = 10µm. **C-D**. Quantification of the number of DDX3X (red) and DDX3X-G3BP1 (yellow) granules per cell from ≥ 30 cells across three independent experiments to monitor the formation of persistent DDX3X-SGs due to DDX3X syndrome mutations in HeLa cells. ****p < 0.0001, ***p = 0.0002, **p = 0.0073 (WT vs WT-WO), **p = 0.0046 (R475G vs R475G-WO), ns = not significant (DDX3X granules/cell), ****p < 0.0001, ***p = 0.0001, **p = 0.0096 (G3BP1-DDX3X granules/cell) (one-way ANOVA test). Data shown are mean ± SEM. **E**. Representative confocal microscopy images of N2a cells expressing WT or mutant DDX3X-mCherry showing SGs upon SA treatment and after wash-off (recovery), immunostained for G3BP1 (green) and DAPI (blue). Scale bar = 10µm. **F-G**. Quantification of the number of DDX3X (red) and DDX3X-G3BP1 (yellow) granules per cell from ≥ 50 cells across three independent experiments to monitor the formation of persistent DDX3X-SGs due to DDX3X syndrome mutations in N2a cells., ***p = 0.0006, **p = 0.0033 (WT vs WT-WO), **p = 0.0020 (F182V vs F182V-WO), ns = non-significant (DDX3X granules/cell), ****p < 0.0001, ***p = 0.0003 (G3BP1-DDX3X granules/cell) (One-way ANOVA test). Data shown are mean ± SEM.

Our observations showing persistent SGs in DDX3X mutation-expressing cells prompted us to test whether DDX3X syndrome mutations alter the liquid-like properties of SGs. To understand this, we performed live-cell imaging of SA-treated cells and subjected them to fluorescence recovery after photobleaching (FRAP) to analyze the liquid/solid-like properties of the SGs. As expected, the DDX3X fluorescence signal of SGs in WT DDX3X expressing HeLa cells immediately recovered (within 30S) after photobleaching, indicating that the WT-DDX3X SGs show liquid-like condensate behavior **(Figure 3A-B)**. Interestingly, DDX3X fluorescence of SGs in HeLa cells expressing F182V, I190S, T198P, T204P, T323I, R480G, L556S, and L559H mutants did not recover after photobleaching, although T323I and L559H mutant DDX3X-SGs showed marginal recovery after photobleaching **(Figure 3A-B).** This suggests that DDX3X missense mutants promote solid-like SG formation. Furthermore, we performed FRAP experiments in N2a cells and found that DDX3X syndrome mutations, F182V, I190S, T198P, T323I, L556S, and L559H, led to the formation of solid-like SGs, which did not show fluorescence recovery after photobleaching. However, unlike in HeLa cells, the R480G mutant of DDX3X, spanning the helicase domain, did not recover after photobleaching **(Figure 3C-D).** Overall, these observations indicate that DDX3X syndrome mutations in close proximity to the NTE and in the CTE regions trigger the formation of persistent solid-like SGs.

**Figure 3.**
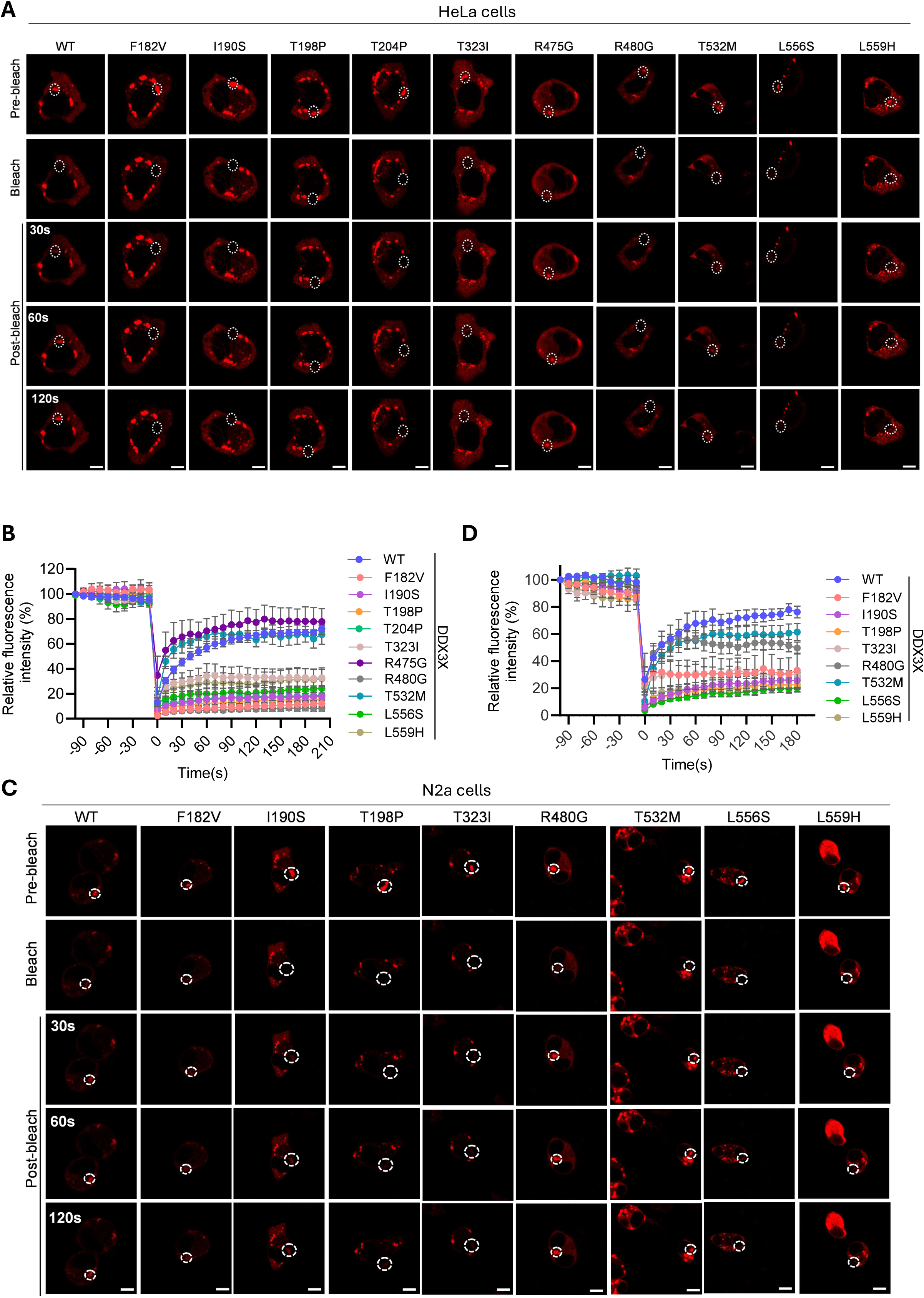
Specific DDX3X syndrome mutations confer solid-like properties to DDX3X-stress granules. **A**. Real-time time-lapse confocal images of HeLa cells expressing WT or mutant DDX3X-mCherry and subjected to SA (200µM) stress for 2 hours. A single SG (indicated by white dashed circle, as the region of interest) was photobleached and fluorescence recovery was recorded over 180s using confocal microscopy. **B**. The relative fluorescence intensity of mCherry (tagged to DDX3X) in SGs in HeLa cells before and after photobleaching as a function of time. Curves are representative of four independent experiments. **C.** Real-time time-lapse confocal images of N2a cells expressing WT or mutant DDX3X-mCherry and subjected to SA (100µM) stress for 2 hours. A single SG (indicated by a dashed circle, as the ROI) was photobleached, and fluorescence recovery was recorded over 150s using confocal microscopy. **D.** The relative fluorescence intensity of mCherry (tagged to DDX3X) in SGs in N2a cells before and after photobleaching as a function of time. Curves are representative of five independent experiments.

### DDX3X mutant-induced persistent granules do not alter translational initiation but promote neuronal lytic cell death

DDX3X promotes translation initiation of selective mRNA subsets, which have structurally complex 5’-UTRs, with global translation largely unaffected at basal conditions ^2; 33; 34^. To test whether the persistent granules formed by DDX3X syndrome mutations affect the global translation and the translation of DDX3X-dependent mRNAs, we performed a polysome profiling assay of N2a cell lysates expressing DDX3X mutants **(Figure 4A-B).** In particular, we focused on NTE proximity (F182V, I190S, and T198P) and CTE (L556S and L559H) mutations in DDX3X and found that the persistent granules formed by these mutants did not exhibit any significant global translational defects compared to WT-DDX3X-expressing cells **(Figure 4B).** To check whether these mutants affect the translation of DDX3X-specific mRNAs, we isolated RNA from pooled polysome fractions and subjected them to RT-qPCR to quantify the DDX3X-specific mRNA fractions. We selected several mRNAs that are dependent on the DDX3X mediated translation initiation (EIF3I, EIF4G2, eIF4E, ATF4, RAC1, TOPBP1, RPL36A, RPL13, STAT1, ODC1, GNB2 and DDX3X). The RT-qPCR data indicate that these DDX3X mutations did not reduce the expression of the DDX3X-specific mRNAs in general, suggesting that these mutations do not affect translation. Intriguingly, L559H, L556S and F182V appeared to increase the translation of some of the DDX3X-dependent mRNAs **(Figure 4C).** These observations indicate that DDX3X syndrome mutations-induced persistent granules do not significantly alter the global translation and might impact the translation of a few DDX3X-dependent mRNAs.

**Figure 4.**
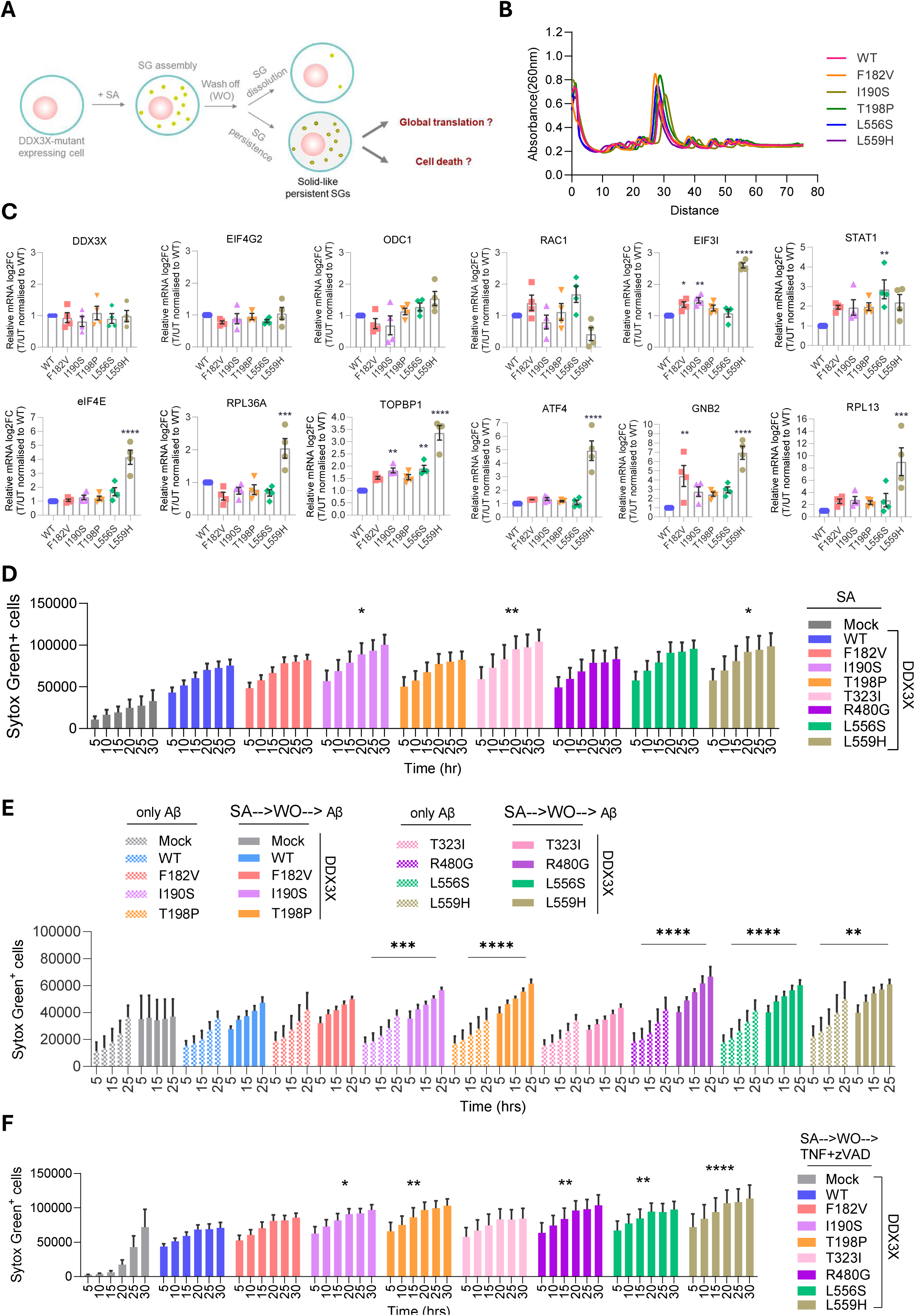
DDX3X missense variants preserve global translation but promote neuronal cytotoxicity. **A.** Schematic of the assay protocol to assess the effect of persistent SGs on global translation and cell death. **B.** Representative polysome profiles (A260 absorbance) of DDX3X syndrome mutants expressing N2a cell extracts treated with SA (100uM) followed by wash-off and recovery in SA-free media, indicating sedimentation (through 10%-50% sucrose gradient) of 40S & 60S ribosomal subunits, 80S monosomes and polysomes (n=3). **C.** Quantification of DDX3X-dependent mRNAs in the polysome fractions plotted as relative log2-fold change ratio of translated/untranslated fraction using gene-specific primers normalised to GAPDH (internal control). ****p < 0.0001, ***p = 0.0007 (WT vs L559H, RPL36A gene), ***p = 0.0002 (WT vs L559H, RPL13 gene), **p = 0.0034 (WT vs I190S, EIF3I gene), **p = 0.0024 (WT vs L556S, STAT1 gene), **p = 0.0035 (WT vs I190S, TOPBP1 gene), **p = 0.0016 (WT vs L556S, TOPBP1), **p = 0.00365 (WT vs F182V, GNB2 gene), *p = 0.0347 (One-way ANOVA). Data shown are mean ± SEM. **D.** Real-time cell death analysis by Sytox green staining of N2a cells expressing WT or mutant DDX3X-mCherry constructs treated with SA **p = 0.0079, *p = 0.0432 (WT vs I190S), *p = 0.0306 (WT vs L559H) (two-way ANOVA test). Data shown are mean ± SEM. **E.** Real-time cell death analysis by Sytox green staining of N2a cells expressing WT or mutant DDX3X constructs treated with Aβ peptide with or without SA treatment and wash off. ****p < 0.0001, ***p = 0.0004, **p = 0.0068 (two-way ANOVA test). Data shown are mean ± SEM. **F.** Cell death measurement by Sytox green staining of N2a cells expressing WT or mutant DDX3X treated with SA, followed by TNF and zVAD treatment. ****p < 0.0001, **p = 0.0020 (WT vs T198P), **p = 0.0038 (WT vs R480G), **p = 0.0053 (WT vs L556S), *p = 0.0182 (two-way ANOVA test). Data shown are mean ± SEM.

Persistent granules deregulate cellular homeostasis and might promote cytotoxicity ^35;36^. Persistent biological aggregates are known to promote neuronal cytotoxicity and neurodegeneration ^32; 37; 38^. Additionally, Aβ aggregates trigger the activation of lytic inflammatory cell death, leading to neuroinflammation ^39–42^. To understand whether persistent SGs formed by DDX3X syndrome mutants promote lytic form of cell death activation, N2a cells were subjected to persistent granule formation followed by treatment with the pre-formed fibrils of Aβ_1-42_ peptide. Real-time analysis of N2a cell death using Sytox Green staining showed that WT-DDX3X or its mutants’ expression enhanced N2a cell death compared to mock-transfected cells, and no apparent differences were observed in DDX3X-mutant-induced cell death compared to WT-DDX3X **(Figure S3A).** The I190S (NTE spanning), T323I (helicase domain), and L559H (CTE spanning) mutants of DDX3X showed a marginal but significant increase in N2a cell death after SA treatment **(Figure 4D).** Also, we observed no significant alteration in cell death levels in Aβ_1-42_ peptide-treated WT-DDX3X expressing N2a cells with or without prior SA treatment. However, Aβ_1-42_ peptide-treatment significantly augmented lytic cell death in N2a cells harboring persistent granules due to the expression of I190S, T198P, R480G, L556S, and L559H DDX3X mutants **(Figure 4E).** This suggests increased cytotoxicity in N2a cells showing persistent granules due to these DDX3X syndrome mutations.

To further establish the role of DDX3X-mutant-induced persistent granules in cellular cytotoxicity, we treated DDX3X-mutant expressing N2a cells with programmed necrosis trigger (TNF+zVAD) after persistent granule formation. We observed no apparent differences in cell death levels of DDX3X-mutant-expressing N2a cells compared to WT-DDX3X-expressing cells after TNF+zVAD treatment, without persistent granules **(Figure S3B)**. DDX3X mutants, I190S, T198P, R480G, L556S, and L559H, showed significantly increased cell death levels after persistent granule formation followed by TNF+zVAD treatment **(Figure 4F).** Thus, these observations suggest that DDX3X syndrome mutations near the NTE and in the CTE regions form persistent solid-like granules promote neuronal cytotoxicity, particularly through the lytic form of cell death. Although the effect of helicase domain spanning R480G mutation of DDX3X show cell type specific persistent SG formation, it still promoted cellular cytotoxicity, suggesting a possible persistent SG-independent function.

### DDX3X mutations show distinct molecular aggregation behaviors in vitro

Our observations demonstrate that NTE and CTE mutations in DDX3X syndrome lead to persistent SGs and cellular cytotoxicity without altering the cellular translation functions. We aimed to understand how these mutations alter the DDX3X structure and LLPS properties to form persistent SGs. To investigate this, we purified the functionally active DDX3X along with NTE and CTE mutants that preferentially promoted persistent granules (F182V, I190S, T198P, L556S, and L559H), which retain a significant portion of the N & C-termini intrinsically disordered regions (residues 132-607) **(Figure 5A-B)**. The DDX3X WT and mutant constructs were tagged with MBP to improve the solubility of the protein during purification **(Figure 5A-C).** To test the effect of DDX3X syndrome mutations on solubility and stability, we performed a precipitation assay in which the MBP-tagged proteins were incubated overnight with TeV protease, and the soluble and insoluble fractions were subjected to SDS-PAGE. The DDX3X mutations did not significantly alter the solubility of the DDX3X upon MBP cleavage **(Figure 5D)**. However, the L556S mutation in DDX3X appeared to reduce protein solubility and led to increased DDX3X in the insoluble fraction **(Figure 5D).** Monitoring the folding and secondary structures of the purified DDX3X proteins, using CD spectroscopy, showed that CTE DDX3X mutations, L556S and L559H showed a hypsochromic shift compared to WT-DDX3X and other mutants, indicating altered protein secondary structures **(Figure 5E).** Further staining with ThT fluorescent dye, which indicates the misfolding and the accessibility of the hydrophobic patches, showed that L556S mutation led to significantly higher binding of ThT at 0 h of incubation. However, all the mutant proteins showed ThT binding after 24 h of incubation **(Figure 5F).** Overall, the C-terminal mutants of the DDX3X appeared to alter the stability and solubility of the DDX3X.

**Figure 5.**
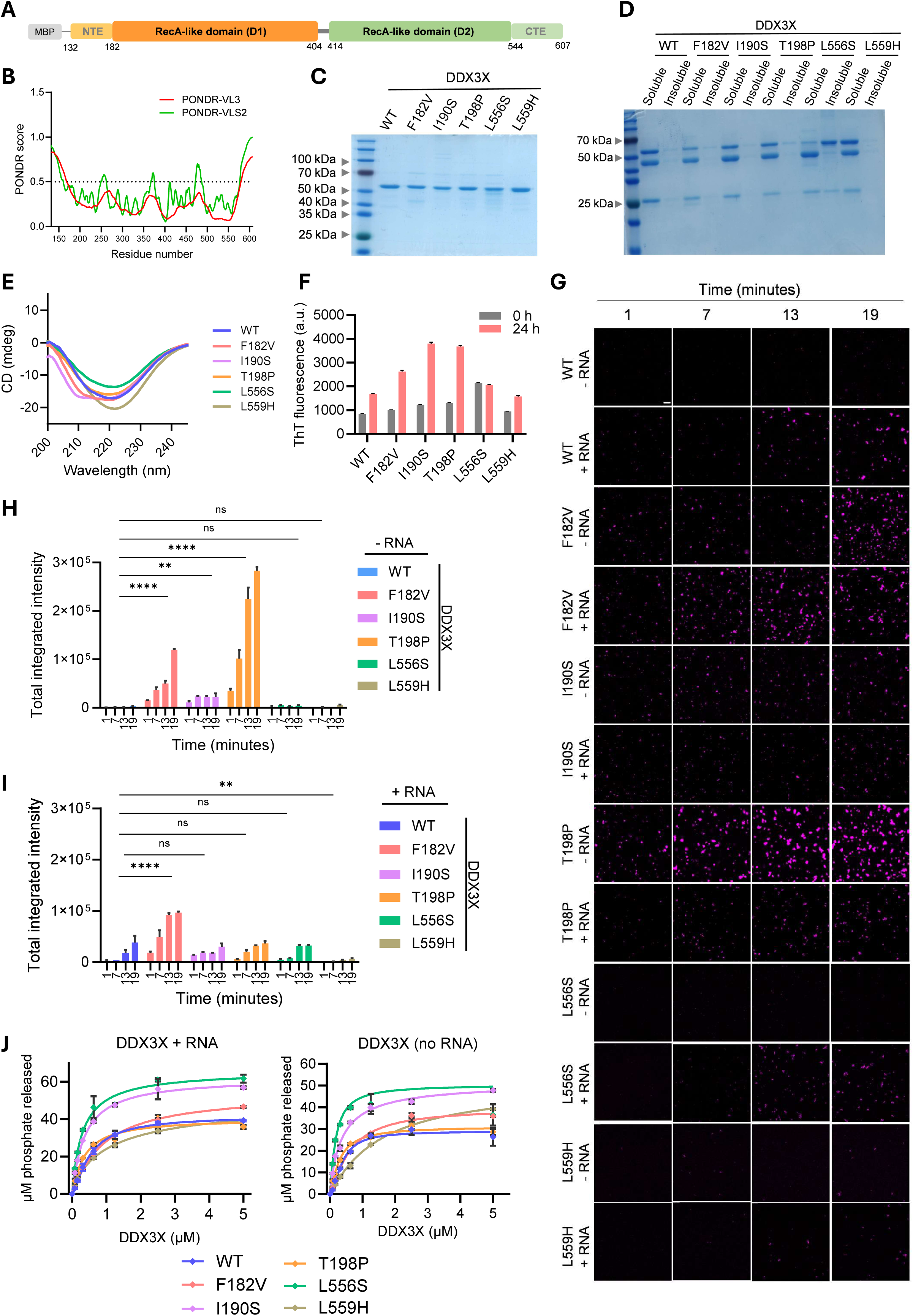
Biophysical characterization of NTE and CTE mutations in DDX3X syndrome reveals an ATPase-independent role in persistent granule formation. **A.** Schematic representation of the MBP-tagged DDX3X construct used for purifying recombinant DDX3X. **B.** Overlay of the predicted disordered regions in DDX3X showing PONDR-VL3 (red) & PONDR VSL2 (green) disordered propensity plots. **C.** SDS-PAGE visualization showing the purified WT and DDX3X mutants by Ni-NTA affinity chromatography. **D.** SDS-PAGE gel showing partitioning of the DDX3X WT and mutations after TeV protease-mediated cleavage of the MBP-tag. Following cleavage, samples were separated into soluble and insoluble fractions. Bands corresponding to the MBP tag (∼100kDa), cleaved DDX3X protein (∼50kDa), and TeV protease (∼25kDa) are shown. **E.** CD spectra profiling of purified WT and mutant DDX3X proteins to monitor secondary structure patterns. **F.** ThT fluorescence intensity at 490 nm at 0 and 24 hours of incubation indicates structural changes and protein-misfolding in the DDX3X protein over time. **G.** In vitro phase separation of 10uM purified and Cy5-labelled WT and DDX3X mutants in the presence and absence of 50ng RNA. **H-I**. Quantification of the total integrated intensity of DDX3X condensation in the absence and presence of RNA, as seen in Panel-G. ****p < 0.0001, **p = 0.0031 (WT vs I190S, -RNA), **p = 0.0024 (WT vs L559H, +RNA), ns = non-significant (Two-way ANOVA test). Data shown are mean ± SEM. **J**. ATPase activity of WT and DDX3X mutants measured after a 30-minute reaction in the presence of 0.5 mg ATP and varying concentrations of protein from 0 to 5 μM in the presence or absence of presence of 100 ng purified total cellular RNA.

To understand how the persistent granule-forming DDX3X mutations could be affecting the protein condensation, we performed in vitro condensate formation (a measure of phase separation) of purified proteins in the presence and absence of RNA. DDX3X proteins were labeled with the Cy5 fluorophore for real-time monitoring of condensation. Purified WT-DDX3X protein formed condensates in the presence of RNA. Unlike WT-DDX3X protein, F182V, I190S, and T198P mutants (N-terminal mutants) showed condensate formation in the absence of RNA **(Figure 5G).** Imaging-based total integrated fluorescence intensity measurements indicate that mixing RNA with the F182V and I190S mutant proteins did not increase condensate formation, and the T198P mutant showed a reduction in condensate formation after adding RNA. Notably, L556S and L559H mutants (C-terminal mutations) did not show condensation in the absence of RNA and showed lesser condensate formation than WT-DDX3X in the presence of RNA **(Figure 5H-I).** These observations indicate that N-terminal DDX3X syndrome mutations promoted homomeric aggregation and DDX3X condensation, whereas C-terminal mutations reduced DDX3X condensation.

Although the N and C-terminal DDX3X mutants formed persistent solid-like SGs in cells, the NTE proximity mutations augmented DDX3X condensation in purified protein conditions (particularly, F182V and T198P mutants), but not the CTE mutations. Thus, N and C-terminal mutations of DDX3X drive the assembly of persistent granules in distinct mechanisms. The ATPase activity of the DEAD-box proteins is essential for the liquid-like properties, and loss of ATPase activity might promote solid-like condensate formation ^14; 43^. We examined the ATPase activity of DDX3X mutants using phosphate release assays to understand whether NTE and CTE mutants of DDX3X blunt the ATPase activity, leading to less dynamic granule formation. Except the L559H, which showed slow phosphate release at lower protein concentration, none of the DDX3X mutants showed reduced ATPase activity. I190S, and L556S showed a detectable increase in ATPase activity of the DDX3X **(Figure 5J).** These observations suggest that the N and C-terminal mutations of DDX3X exhibit ATPase activity-independent, mutation-specific regulation of DDX3X solid-like condensate formation. Recent studies showed that ATPase activity of DDX proteins does not necessarily correlate with the liquid-like condensate properties ^29; 44^, further indicating an ATPase-independent mechanism of N-and C-terminal DDX3X mutations.

### DDX3X syndrome mutations confer conformational bias of DDX3X to its open conformation and the rigidity of the DDX3X-RNA complexes to promote persistent granule formation

We sought to perform molecular dynamics simulations to understand how NTE and CTE mutations distinctly impact DDX3X structure dynamics and RNA binding. The structure of the DDX3X (PDB: 6O5F) shows a helicase core composed of two RecA-like domains (D1 and D2), a significant part of the N and C-termini, and a few missing residue coordinates in the helicase core. We modeled and refined the missing loop regions of the DDX3X structure, and the modeled loop regions did not alter the core DDX3X 3D-structure **(Figure 6A).** Using this refined DDX3X structure, we introduced the DDX3X mutations and generated the refined DDX3X mutant models. To monitor the impact of DDX3X mutations on DDX3X structural dynamics and its interactions with RNA, we performed MD simulations of WT-DDX3X and the mutants, both with and without RNA. The D1 and D2 domains of DDX3X remain in an open conformation to enable accessibility of the RNA-binding interface **(Figure 6C).**

**Figure 6.**
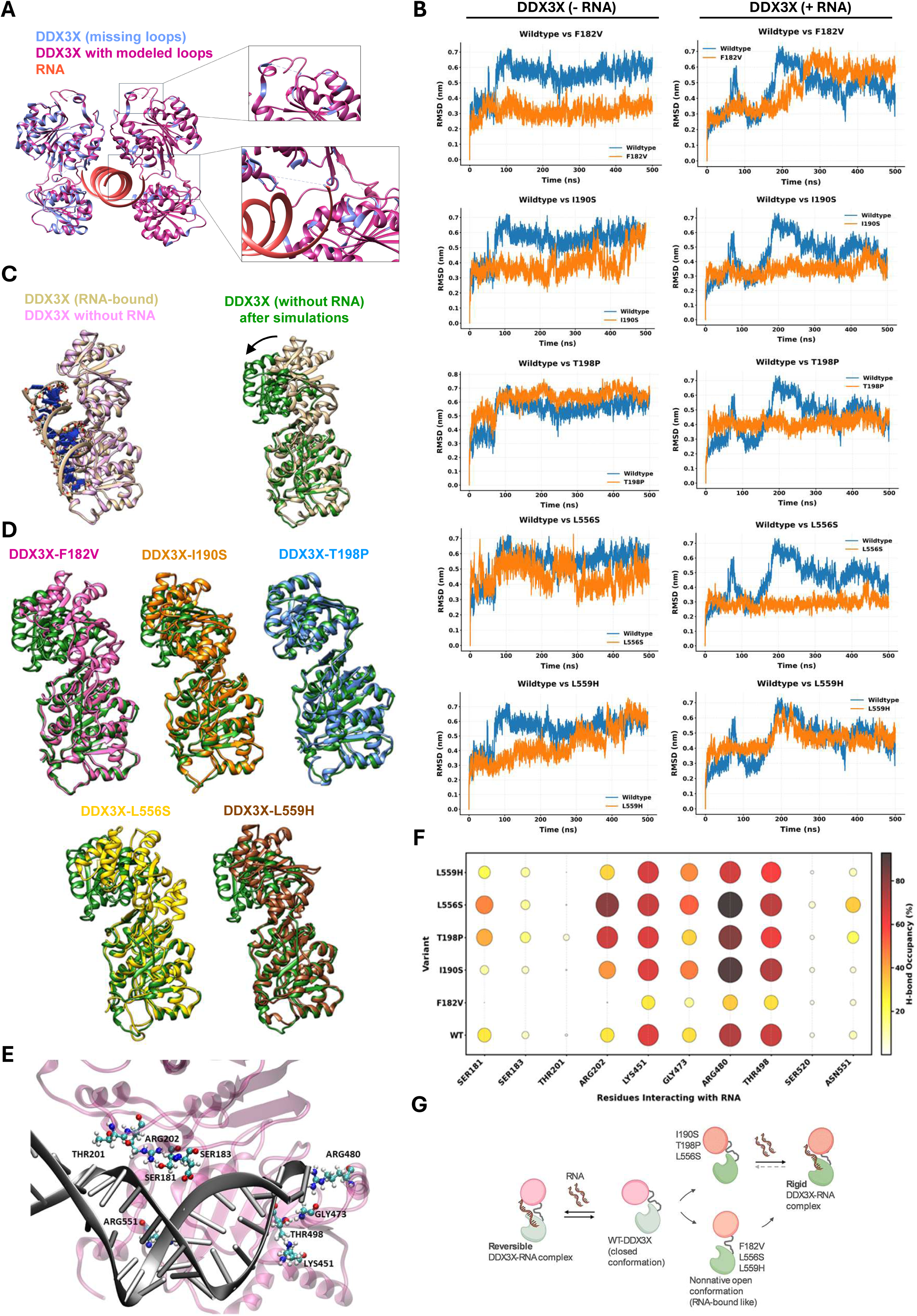
Molecular dynamics (MD) simulations of DDX3X syndrome mutations reveal distinct DDX3X-RNA conformational preferences. **A.** Structural superposition of the DDX3X (cyan) and the modelled DDX3X structure illustrating the reconstructed loop regions (pink). **B.** MD simulations-based Root Mean Square Deviation (RMSD) of WT and DDX3X either with RNA or without RNA, showing the comparison of the WT-DDX3X trajectory (blue) against the mutant trajectories (orange). **C.** Superimposition of the WT-DDX3X in its RNA-bound and RNA-unbound structures in conformation prior to the simulations, indicating both starting conformations are structurally similar prior to the MD simulation. **D**. Structural comparison of the WT-DDX3X protein before and after simulations. The structure in green represents the closed conformation obtained after simulations, overlaid on the initial (beige) structure, which is in the RNA-bound open conformation. **D**. Structural comparison of the WT-DDX3X (closed conformation, in green) with the indicated DDX3X-mutant structures. **E**. Structural representation of DDX3X-RNA interactions highlighting key residues involved in RNA recognition. The DDX3X is in pink, and the RNA is in grey. The RNA-binding residues are displayed as ball-and-stick models and labelled accordingly. **F**. Bubble heat map plots showing hydrogen-bond occupancy between RNA and the interacting residues of DDX3X during the MD simulation of WT and the indicated DDX3X mutations. Bubble size and color intensity indicate the percentage occupancy of hydrogen bonds throughout the trajectory. **G**. Schematic showing the effect of specific DDX3X syndrome mutations on DDX3X conformational alterations favoring nonnative open conformation (RNA-bound like) and the stabilization of rigid DDX3X-RNA conformations.

MD simulations demonstrate that N-terminal (F182V, I190S, T198P) and C-terminal (L556S, L559H) mutations of DDX3X perturb its conformation at distinct stages **(Figure 6B)**. WT-DDX3X (without RNA) transitioned to the closed state due to the movement of D2 into proximity to D1, thereby masking the RNA binding interface **(Figure 6C).** The C-terminal mutations, L556S and L559H, and the N-terminal F182V restrict the movement of the D2 domain and remain in the open state, locking the DDX3X structure in a conformation suitable for RNA binding and limiting DDX3X transition to the closed state **(Figure 6D)**.

In contrast, N-terminal mutations, I190S and T198P, and the C-terminal L556S mainly affected the RNA-bound state of the DDX3X by restricting the conformational sampling of the DDX3X-RNA complex **(Figure 6B & Figure S4A).** This likely locks the DDX3X-RNA state and impairs the conformational transitions, possibly essential for RNA unwinding. Notably, RNA-DDX3X interaction analysis indicated that none of these DDX3X mutants (except F182V) alter the overall RNA-binding interface and increase the RNA interactions compared to the WT-DDX3X **(Figure 6E-F).** Thus, MD simulation analysis indicates that these mutants drive the persistent DDX3X aggregation either by locking DDX3X in an open (RNA-bound like) conformation or by increasing the rigidity of the DDX3X-RNA complex. To validate these observations, we performed simulations with the R475G and R480G mutants of DDX3X (which did not form persistent DDX3X-SGs) and found that these mutations do not alter the DDX3X conformations in either the RNA-bound or unbound states **(Figure S4B-C)**.

### DDX3X syndrome mutations in CTE promote Aβ aggregation and amyloid-like granules

Pathological protein aggregates disrupt neuronal cell physiology, driving neurodegeneration and neuroinflammation ^25; 27; 32^. Particularly, amyloid-β aggregates are associated with several neurodevelopmental disorders ^24–27^. The biochemical and MD-simulation studies indicate that, unlike N-terminal mutations, C-terminal mutations in DDX3X (L556S and L559H) alter protein stability and folding. This may result in partially folded DDX3X conformations that can predispose the protein to cross-seeding other neuronal aggregates like amyloid-β (Aβ). To test whether C-terminal DDX3X mutations confer aggregate cross-seeding ability, we performed Aβ peptide aggregation in the presence of WT-DDX3X and its mutant proteins. The purified DDX3X protein and its mutants were allowed to form aggregates and sonicated to generate small seeds suitable for cross-seeding the Aβ peptide **(Figure 7A).** As a control, the sonicated Aβ_1-42_ peptide aggregates were used, which promote Aβ peptide aggregation. Thioflavin T (ThT) fluorescent dye, which stains β-sheet–rich amyloid-like structure, was used to monitor Aβ peptide aggregation kinetics, and the T_1/2_ values were measured, which indicates the amount of time at which half of the Aβ peptide is aggregated by the DDX3X/Aβ_1-42_ seeds. Notably, only L556S and L559H mutations showed a considerable decrease in the T1/2 value of Aβ aggregation (29% and 36% reduction, respectively), similar to the effect shown by the Aβ_1-42_ seed (decrease in T_1/2_ by 32.8%) **(Figure 7B).** Only DDX3X seeds did not show any apparent increase in ThT fluorescence, suggesting the absence of amyloid-like aggregation **(Figure S5)**. This indicates that the L556S and L559H mutations in DDX3X promote the cross-seeding of Aβ_1-42_ aggregates.

**Figure 7.**
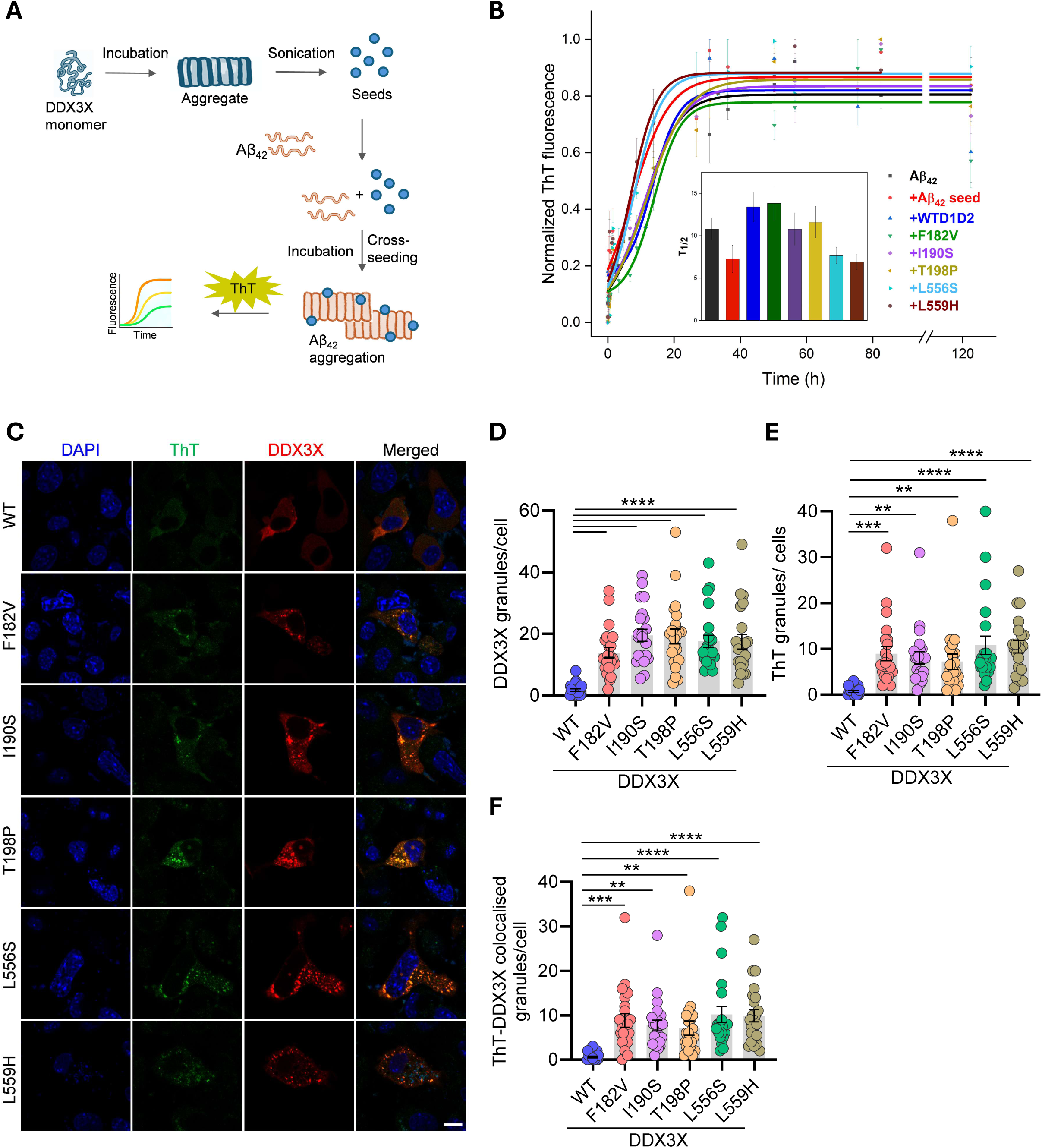
C-terminal DDX3X syndrome variants cross-seed Aβ_1-42_ aggregates. **A**. Schematic showing the protocol used for the cross-seeding kinetics of Aβ_1-42_ aggregation by DDX3X seeds. **B**. Boltzmann fitting of the cross-seeding experiment depicting the co-aggregation kinetics of Aβ_1-42_ when cross-seeded with sonicated seeds of DDX3X variants. **C**. Representative confocal images of N2a cells expressing WT or mutant DDX3X-mCherry, subjected to SA stress (100µM), wash-off, and recovery, followed by ThT staining and immunostained with DAPI (blue). ThT fluorescence is represented in green. Scale bar= 10µm. **D-F**. Quantification of numbers of DDX3X (red), ThT stained (green) and ThT-DDX3X (yellow) granules per cell from ≥ 20 cells across three independent experiments exhibiting DDX3X colocalization with ThT granules in N2a cells. ****p < 0.0001 (DDX3X granules/cell), ****p < 0.0001, ***p = 0.0005, **p = 0.0022 (WT vs I190S), **p = 0.0080 (WT vs T198P) (ThT granules/cells) ****p < 0.0001, ***p = 0.0003, **p = 0.0021 (WT vs I190S), **p = 0.0057 (WT vs T198P) (ThT-DDX3X colocalised granules/cells) (One-way ANOVA test). Data shown are mean ± SEM.

Given that the L556S and L559H mutants of DDX3X appeared to promote amyloid-β cross-seeding, we further tested whether these mutants confer amyloid-like properties to the persistent stress granules in cells by ThT staining of cells **(Figure 7C).** As expected, the DDX3X mutants showed persistent DDX3X-granules in N2a cells compared to WT-DDX3X-expressing cells. All the DDX3X mutants expressing cells, but not the WT-DDX3X, showed DDX3X-granules stained with ThT, although fewer in number than the DDX3X-stained granules. Notably, the C-terminal mutants, L556S and L559H, showed a higher number of ThT granules than the N-terminal mutants **(Figures 7D-F)**. These observations indicate that DDX3X-syndrome mutants that induce persistent granules exhibit amyloid-like properties, and that C-terminal DDX3X mutants promote amyloid aggregation.

## DISCUSSION

DDX3X is a highly conserved DEAD-box RNA helicase regulating critical cell fate choices, and mutations in this protein potentially drive profound cellular and organismal dysfunction ^3; 9; 16; 45^. Individuals with DDX3X syndrome, resulting from mutations in the DDX3X, exhibit a broad spectrum of neurodevelopmental abnormalities ^17; 20; 21; 46^, underscoring the crucial role of this protein in neuronal homeostasis. While the phenotypic landscape of DDX3X-associated disorders has been well documented, the molecular mechanisms connecting individual mutations to diverse clinical outcomes remain largely unresolved. This study provides evidence that missense mutations of DDX3X syndrome patients, in both the N- and C-terminal regions, converge on a common pathological outcome: the formation of aberrant, persistent, solid-like stress granules that alter cell fate toward lytic cell death. Notably, although these mutations display distinct aggregation behaviors in vitro, they consistently promote persistent solid-like granules in cells. These observations suggest that altered molecular interactions in the stress granules, rather than initial assembly kinetics, may be the critical determinant of DDX3X syndrome mutations induced cellular toxicity. These persistent granules likely act as seeds for pathological protein aggregation, revealing a potential link between DDX3X dysfunction and broader neurodegenerative processes. Previous work has implicated DDX3X in neuronal progenitor development through its regulation of RNA granules ^17; 46^, although no mechanism has been identified. Our findings demonstrate that DDX3X syndrome–associated missense mutations significantly alter the material properties of phase-separated granules, leading to a transition from dynamic, liquid-like assemblies to rigid, solid-like structures. This shift in granule state appears to have profound impact on cell fate decisions, promoting neuroinflammatory signalling. Importantly, several of the mutations examined here have also been reported in cancer^10; 16; 23^, raising the intriguing possibility that granule persistence–mediated lytic cell death may represent a shared pathogenic mechanism across neurodevelopmental disorders and cancer progression.

Mechanistic insight into how DDX3X mutations promote persistent granules emerges from our structure-based simulations of DDX3X–RNA complexes. These analyses reveal that pathogenic mutations impose distinct conformational preferences on DDX3X, likely predisposing the protein to aberrant assembly states. N-terminal mutations, mainly I190S and T198P, preserve the compact conformation of RNA-free DDX3X, however, they markedly restrict conformational flexibility of RNA-bound DDX3X. This leads to unusually rigid RNA–protein complexes that favor stable intermolecular interactions and sustained solid-like granule formation. In contrast, the F182V mutation induces comparatively modest conformational changes, consistent with its milder biophysical effects. C-terminal mutations, including L556S and L559H, exhibit a distinct impact on DDX3X folding and conformation. These variants appear to trap DDX3X in an open, RNA-bound–like conformation, interfering with normal allosteric transitions that precede RNA engagement. Such alterations likely reflect broader folding defects and may explain the enhanced propensity of these mutants to cross-seed other protein aggregates, including amyloid-like assemblies. A recent study indicates that the L556S confers purified DDX3X aggregation, further corroborating our observations ^47^. Together, these findings highlight how spatially distinct mutations can perturb distinct steps of the DDX3X conformational cycle while converging on a shared pathological consequence.

Overall, our study establishes a unifying framework in which DDX3X syndrome mutations reprogram the physical behavior of DDX3X-condensates to form pathological persistent granules causing neuronal cell death. By connecting DDX3X mutation-specific conformational changes to granule persistence and inflammatory cell death, this work provides a mechanistic foundation for understanding how DDX3X dysfunction contributes to neurodevelopmental disease and cancer, and points toward granule material properties as a potential therapeutic target.

## METHODS

### Cell culture, transfection and treatment

HeLa and Neuro-2a were cultured in DMEM supplemented with 10% FBS and 1% antibacterial-antimycotic in a humified incubator with 5% C0_2_ at 37^0^C. HeLa cells were transfected using Polyethylamine, linear-25,000 (Polysciences, Inc) and Neuro-2a cells were transfected with Lipofectamine 2000 (Invitrogen). For the formation of granules, equal amounts of plasmid were transfected for each construct followed by Sodium(-meta) arsenite (Sigma Aldrich, S7400) treatment after 24 hours and cells were then fixed after two hours following Sodium(-meta) arsenite stress.

### Constructs

Full length DDX3X-mCherry construct (pReceiver-M56-DDX3X-mCherry ) was procured from GeneCopoeia. The WT and mutant DDX3X-mCherry plasmids were generated by subcloning into pLVX-EF1alpha-GFP-IRES-Puro vector backbone, a kind gift from Prof. Nevan Krogan’s lab (University of California, San Francisco) using EcoRI and BamHI restriction sites. Point mutations on DDX3X (all mutations) were generated using the PrimeStar Max (TakaraBio) mediated site-directed mutagenesis and overlap-extension PCR method.

The human DDX3X core (residues 132-662; PDB:6O5F) gene was synthesised by GeneArt and cloned into pMAL-C2X vector backbone (Addgene #75286) using EcoRI and SbfI restriction sites. All the constructs consist of N-term 6X-His tag with the TEV protease cleavage site between the MBP-tag and gene of interest. Each of the DDX3X mutations, were introduced by site-directed mutagenesis. All plasmids were verified by restriction digestion and Sanger’s sequencing method.

### Immunofluorescence

HeLa and Neuro-2a were seeded in 8-chambered glass slides and cultured overnight. The WT and mutants of DDX3X were transfected using PEI (Polysciences, Inc) for HeLa and Lipofectamine 2000 (Invitrogen) for Neuro-2a. After 24 hours, cells were treated with sodium (-meta) arsenite (Sigma Aldrich, S7400) for 2 hours and then fixed with 4%PFA (paraformaldehyde) at room temperature (RT) for 15 minutes followed by washing of the cells thrice with PBS and permeabilized with 0.1% Triton-X-100 in PBS for 10 minutes at RT. After washing the cells twice with PBS, the cells were blocked with 3% BSA (GBiosciences) in PBS for 1hr at RT. For stress granule imaging, anti-G3BP1 (Invitrogen), was diluted in 1:200 ratio and kept overnight at 40C. Following this, cells were washed four times with PBS and then treated with anti-rabbit Alexa-fluor 488 (Invitrogen) secondary antibody diluted at a ratio of 1:500 for 1 hour at room temperature. After washing the cells four times with PBS, the cells were counterstained with 0.5ug/ml DAPI for 1-2 minutes and then replaced with PBS. All images were acquired using Olympus FV 300 confocal microscope. Quantification of the puncta per cell was done using ImageJ and the results were plotted using GraphPad Prism software.

For wash-off assay, HeLa and Neuro-2a were seeded in 8-chambered glass slides and cultured overnight followed by transfection of WT and mutant DDX3X. After 24 hours, cells were treated with sodium (-meta) arsenite for 2 hours and then replaced with fresh media. Fixation of the wash-off set was done using 4%PFA (paraformaldehyde) at room temperature (RT) for 15 minutes followed by washing of the cells thrice with PBS and permeabilized with 0.1% Triton-X-100 in PBS for 10 minutes at RT. After washing the cells twice with PBS, the cells were blocked with 3% BSA (GBiosciences) in PBS for 1hr at RT. For stress granule imaging, anti-G3BP1 (Invitrogen), was diluted in 1:200 ratio and kept overnight at 40C. Following this, cells were washed four times with PBS and then treated with anti-rabbit Alexa-fluor 488 (Invitrogen) secondary antibody diluted in the ratio of 1:500 for 1 hour at room temperature. After washing the cells four times with PBS, the cells were counterstained with 0.5ug/ml DAPI for 1-2 minutes and then replaced with PBS. All images were acquired using Olympus FV 300 confocal microscope. Quantification of the puncta per cell was done using ImageJ and the results were plotted using GraphPad Prism software.

### Fluorescence recovery after photobleaching (FRAP)

The FRAP assays were conducted using the bleaching module of Olympus FV 300 Confocal microscope. The 561nm laser was used to bleach the mCherry signal.

HeLa and Neuro-2a cells were seeded on 8-chambered glass slide (Cellvis) and the plasmids of WT and mutant DDX3X were transfected into HeLa and Neuro-2a cells (in a separate set of experiment). After 24 hours, the normal DMEM media was replaced with sodium (-meta) arsenite (Sigma Aldrich, S7400) treatment media and taken for imaging after 2hours. To perform a full bleach assay, an entire punctum was selected as an ROI to bleach with 100% laser power and time lapse images were collected afterward. The fluorescence intensity was directly measured using ImageJ software. The values were reported relative to pre-bleaching time points. GraphPad Prism was used to plot the data. The halftime of each replicate was calculated using the following formula:

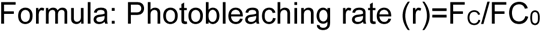

Where, F_C_= unbleached area after photobleaching

FC0= unbleached area before photobleaching

Photobleaching intensity (F)= (F_S_-F_B_)/r

Where, F_S_= bleached area

F_B_= blank (dark) area

### Cell-death assay

Real-time cell death analysis were performed using a two-colour IncuCyte S3 Live-Cell analysis instrument (Sartorius). Neuro-2a cells were seeded in 48-well plates (Eppendorf) and transfected with either WT or DDX3X mutant constructs. 24-hours post transfection cells were treated with sodium (-meta) arsenite treatment for 1.5 hours and then replaced with fresh media (recovery) followed by treatment with Aβ peptide or programmed necrosis triggers (zVAD+TNF). Dead cells were stained with 20nM sytox green (Thermo Fisher Scientific), a cell-impermeable DNA-binding fluorescent dye which rapidly enters dying cells after membrane permeabilization and fluoresces green. The resulting images and the fluorescence signals were analysed using IncuCyte S3 software, which provides sytox-green positive cell count. The data were further plotted using GraphPad Prism 9.0 software.

### Polysome profiling

Neuro-2a cells were seeded in 100mm dishes followed by transfection with WT or DDX3X mutants. The transfected cells were treated with 100uM SA followed by two hours of recovery period. Cells were treated with 100µg/ml cycloheximide (Sigma) for 15 minutes, washed with ice-cold PBS supplemented with 100µg/ml cycloheximide twice and harvested into lysis buffer (5mM Tris-HCL; 2.5mM MgCl_2_; 1.5mM KCl; 0.5%Triton-X; 0.5%sodium deoxycholate; 100µg/ml cycloheximide; 2mM DTT) supplemented with RNase plus inhibitor and Phosphatase and protease inhibitors. Lysates were centrifuged at 4^0^C for 10 min at 13,000xg. Supernatants were weighed and loaded onto 10%-50% sucrose gradients made in gradient buffer and centrifuged in a Beckman SW41 rotor (Beckman-Coulter) at 39000r.p.m for 2h at 4^0^C. Samples were eluted and loaded in the profiler and its OD 254 was measured by Biocomp. The untranslated (till 80S fraction) and translated (polysome fractions) were collected and pooled (Biocomp), followed by RNA isolation from both fractions by TRIzol-chloroform method. Total RNA was also isolated from lysates as input. RNAs were measured by NanoDrop.

### RT-qPCR

RNA was extracted from the collected polysome fractions using Trizol-chloroform method^48^ and reverse transcribed using the cDNA synthesis kit (Applied Biosystem, 43688140), RT-qPCRs were performed using the following primer sets:

DDX3X: 5’-AGGTCGGTGTGAACGGATTTG-3’

5’-TGTAGACCATGTAGTTGAGGTCA-3’

ATF4: 5’-AAGGAGGAAGACACTCCCTCT- 3’; 5’-CAGGTGGGTCATAAGGTTTGG- 3’

eIF4e: 5’-ACCCCTACCACTAATCCCCC- 3’; 5’-CAATCGAAGGTTTGCTTGCCA- 3’

eIF3I: 5’-CAAAGACCCTATCGTCAACGTG- 3’; 5’-GAGCCAGTAAGGACATGCTTG- 3’

eIF4G2: 5’-AGTGCGATTGCAGAAGGGG- 3’; 5’-GTGCTTCGTGCAGGAATCCA- 3’

ODC1: 5’-GACGAGTTTGACTGCCACATC- 3’; 5’-CGCAACATAGAACGCATCCTT- 3’

STAT1: 5’-TCACAGTGGTTCGAGCTTCAG - 3’; 5’-GCAAACGAGACATCATAGGCA-3’

GNB2: 5’-TACACCACTAACAAGGTCCACG - 3’; 5’-CAGATGTTGTCCAAACCCCCA-3’

RAC1: 5’-ATGCAGGCCATCAAGTGTG - 3’; 5’-TAGGAGAGGGGACGCAATCT - 3’

RPL13: 5’-AGCCGGAATGGCATGATACTG - 3’; 5’-TATCTCACTGTAGGGCACCTC-3’

RPL36A: 5’-GTGAACGTGCCTAAAACCCG - 3’; 5’-AGGCTTAGTCTGCCCACCATA- 3’

TOPBP1: 5’-CAGGATTGTTGGTCCTCAAGTG - 3’ ;

5’-CCTGCAATAAGGTGAGTGACTG-3’

GAPDH: 5’-AGGTCGGTGTGAACGGATTT-3’; 5’-TGTAGACCATGTAGTTGAGGTC-3’

### Protein expression and purification

All constructs were transformed into BL21 E.coli cells and positive clones were then expressed in LB broth media induced with 1mM IPTG at 16-18^0^C overnight. Cells containing recombinant protein were harvested by centrifugation at 4000g at 4^0^C for 10 minutes and stored at -80 degrees Celsius. Cell lysis was done by dissolving the pellet in binding buffer (50mM Tris-pH 8, 500mM NaCl, 30mM imidazole and 5% glycerol) supplemented with Bug Buster (1.5ml) and PMSF (1mM). The pellets were then sonicated and further centrifuged at 12,500 rpm for 30 minutes at 4^0^C. After centrifugation, the supernatant was filtered using 0.45 um syringe filter and loaded on a His-trap (5ml) column/Ni-NTA column. Protein was eluted by using a linear gradient of imidazole (0-100%). Eluted fractions were further analysed on 12% SDS-PAGE to check the purity of the samples. All the fractions containing the protein were pooled and desalted in desalting buffer (50mM Tris-pH 8, 200mM NaCl and 5% glycerol) using 26/10 hi-Prep Cytiva desalting column. The MBP tag was removed by incubating the purified proteins with TEV protease at a ratio of 1:25 concentration, overnight at 4 °C. The sample was then filtered to remove any precipitate before running through the Ni-NTA column and the flowthrough containing the DDX3X protein without MBP tag was collected and further concentrated using the AMICON 10kDa centrifugal filters, which was stored in aliquots in storage buffer (50mM Tris-pH 8, 100mM NaCl and 5% glycerol) at -80^0^C.

### Precipitation assay

For removal of the MBP tag, 5µM of the purified proteins (DDX3X WT & mutants) was incubated with TEV protease in the ratio of 1:25, overnight at 4 °C in a buffer containing 50mM Tris-HCl (pH 8) and 200 mM NaCl. The incubated Samples were then centrifuged at 15,000rpm for 10minutes and the soluble and the insoluble fractions were separated. The pellets were then resuspended in the buffer. 10μl of both the soluble and insoluble fractions of each sample, boiled in 1X Laemli’s buffer and were run on the SDS-PAGE.

### ATPase assay

The ATPase assay with the purified WT and mutant DDX3X was performed by using a malachite green Phosphate assay kit (Sigma, MAK-307), as described in the literature^29^. Briefly, in 96-well micro-plate assay varying concentration of WT & mutant DDX3X was pre-incubated with 100ng/µl total RNA, extracted from HEK-293T cells, for 15-20minutes in the reaction buffer (50mM Tris-HCL, pH 8, 100 mM NaCl, 1 mM MgCl2) before the addition of 0.5mM ATP. The reaction was incubated at 25 degrees Celsius for 30 minutes. The reaction was then quenched/terminated by adding 20 ul of malachite green reagent and incubated at room temperature for 30minutes for the development of the colour. The absorbance (green colour) was then measured at 620 nm using TECAN infinite M200 Pro. Values were converted from absorbance units to µM free phosphates using a standard curve generated with the kit’s phosphate standard. Data were plotted using GraphPad Prism. Significance was calculated using an unpaired Student’s two-tailed t-test.

### ThT Fluorescence assay

25μM each of purified DDX3X WT & mutants was incubated at 37°C at 1000rpm in Eppendorf thermomixer. The ThT kinetics were then tracked using 20 μM ThT and 2 μM peptide, which were collected at different time points during incubation. The samples were excited at 440 nm, and the emission was recorded at 490 nm using TECAN Infinite M200 Pro multimode microplate reader. All the readings were recorded in duplicates. The fluorescence intensity was plotted against time, and the data were fitted using Boltzmann fitting to calculate the T1/2 value.

### Aβ_1-42_ Cross-seeding kinetics

The incubated DDX3X samples were bath sonicated for 10 minutes to prepare the seeds. 4 μM of the prepared seeds was used to cross-seed 2mg/ml Aβ1-42 dissolved in 25% DMSO in 20µM phosphate buffer and incubated at 37°C at 1000rpm, with constant shaking in an Eppendorf thermomixer. The ThT kinetics were then tracked using 20μM ThT and 2μM peptide, which were picked at different time points during incubation. The samples were excited at 440 nm and the emission was recorded at 490 nm using TECAN Infinite M200 Pro multimode microplate reader. All the readings were recorded in duplicates. The fluorescence intensity was plotted against the time and the data was fitted using Boltzmann fitting to calculate the T1/2 value.

### CD spectroscopy

The CD spectra of 5μM each of WT & mutant DDX3X in storage buffer was recorded using Jasco J-800 CD spectrometer. The spectra were recorded at 25°C from 195 nm to 245 nm using a quartz cuvette of 1mm path length. Final spectra were corrected by subtracting the baseline for buffer composition in all conditions. The spectra were plotted on GraphPad Prism

### Transmission Electron Microscopy

5µl of the incubated samples were drop cast on 200-mesh carbon-coated copper grids (Sigma-Aldrich). This was followed by washing in nuclease-free distilled water and 4% ammonium molybdate solution. The grids were imaged with Talos F200S TAM by Invitrogen.

### In-vitro droplet assay

9 units of unlabelled protein was mixed with 1 unit of Cy-5 labelled proteins in samples containing RNA, 50ng of total RNA was added. All the samples had 5% PEG and 0.04nM of Sytox green. The total reaction was set for 15μl in assay buffer (50mM Tris-pH 8, 100mM NaCl, 1mM MgCl2). The droplets formed were visualised using Olympus FV 300 confocal microscope. The droplets were quantified using image analysing software FIJI.

### Structural Modelling of DDX3X

The tetrameric structure of DDX3X was modelled using MODELLER (v10.7) based on the experimentally determined crystal structure available in the Protein Data Bank (PDB ID: 6O5F). The deposited structure contained multiple unresolved loop regions, which were reconstructed prior to molecular dynamics (MD) simulations. Loop modeling was performed using MODELLER, which applies comparative modeling based on spatial restraints and molecular dynamics–based optimization.

An initial tetrameric model was generated from the 6O5F template and subsequently subjected to sequential loop refinement. Four missing loop regions were refined individually in a stepwise manner. For each loop region, 25 independent models were generated, and model quality was evaluated using the Discrete Optimized Protein Energy (DOPE) score. The model with the lowest DOPE score was selected and used as the input structure for refinement of the subsequent loop. This iterative process was repeated until all four loop regions were fully refined.

### Modelling of Mutant Structures

Mutant models of DDX3X were generated using SWISS-MODEL in template-based modeling mode. The fully loop-refined wild-type DDX3X tetramer was used as the template for all mutant structures to ensure structural consistency across systems.

### System Preparation and Simulation Parameters

All molecular dynamics systems were prepared using CHARMM-GUI. The protein complexes were solvated in an explicit TIP3P water model within a cubic simulation box, ensuring a minimum distance between the protein surface and box edge. Sodium and chloride ions were added to neutralize the system and to achieve physiological ionic strength. The systems were parameterized using the AMBER force field.

### Molecular Dynamics Simulations

Molecular dynamics simulations were performed using GROMACS (v2025) under periodic boundary conditions. Each system was initially energy minimized using the steepest descent algorithm to remove unfavorable steric contacts. The minimized systems were equilibrated under an NVT ensemble to stabilize the temperature at 300 K, with positional restraints applied to protein heavy atoms. Subsequently, production molecular dynamics simulations were carried out for 500 ns with all restraints removed. All simulations were performed with a 2 fs time step. Long-range electrostatic interactions were calculated using the Particle Mesh Ewald (PME) method. Covalent bonds involving hydrogen atoms were constrained using the LINCS algorithm. The resulting trajectories were used for subsequent structural, dynamic, and energetic analyses.

### Molecular Dynamics Trajectory Analysis

Trajectory analyses were performed using GROMACS analysis tools. Prior to analysis, trajectories were processed to remove periodic boundary effects and aligned to the initial structure using protein backbone atoms. Structural stability of the protein systems was assessed by calculating the root mean square deviation (RMSD) of backbone atoms with respect to the starting structure. Global compactness of the protein complexes was examined using the radius of gyration (Rg). Intermolecular interactions were characterized by computing hydrogen bond occupancy between the protein and bound ligands over the course of the simulations, using standard geometric criteria. Time evolution of interaction patterns was analysed over the production trajectory.

## ACKNOWLEDGEMENTS

We thank the S.K. lab members for their comments on this work. We thank Anantika Chandra, Saloni Gehlot, and Subham Saha for helping with initial experiments to express DDX3X constructs and thank Gayatri Mohanan for discussions on polysome assays. We thank Mahipal Ganji from the Indian Institute of Science for helpful suggestions on in vitro phase separation experiments. We thank Naruhiko Sahara from the National Institutes for Quantum Science and Technology, Japan and Gen Matsumoto from Osaka Metropolitan University, Japan for helpful discussions with the protein aggregation assays. S.K. research work is supported by funding from the Indian Council of Medical Research (ICMR) (2021-14148/CMB/ADHOC-BMS) (IIRPSG-2024-01-01943), the Department of Biotechnology (DBT) (BT/PR50450/MED/12/1044/2023), the Anusandhan National Research Foundation (ANRF) (ANRF/ARG/2025/007263/LS), and Scheme for Transformational and Advanced Research in Sciences (STARS)-MoE (MoE-STARS/STARS-2/2023-0464). P.I.R. thanks the Department of Biotechnology (DBT-BT/PR51975/BMS/85/23/2024), India, for supporting research in his laboratory. P.G. and S.L. acknowledge the Ministry of Education (MoE), India, for the research fellowship.

## AUTHOR CONTRIBUTIONS

S.K. conceptualized the study; P.G., S.K.K., and S.K. designed experiments and methodologies; P.G., and S.K.K. performed cell culture and biochemical experiments, in vitro aggregation assays; S.G. performed DDX3X mutant protein structure prediction and the molecular dynamic simulations; S.L. and P.I.R. planned and performed polysome profiling assays; S.K., P.G., and S.K.K. wrote and edited the manuscript; S.K. supervised the study; All the authors contributed to manuscript editing; S.K. provided the guidance and brought the funding.

## SUPPLEMENTAL FIGURE LEGENDS

**Figure S1.**
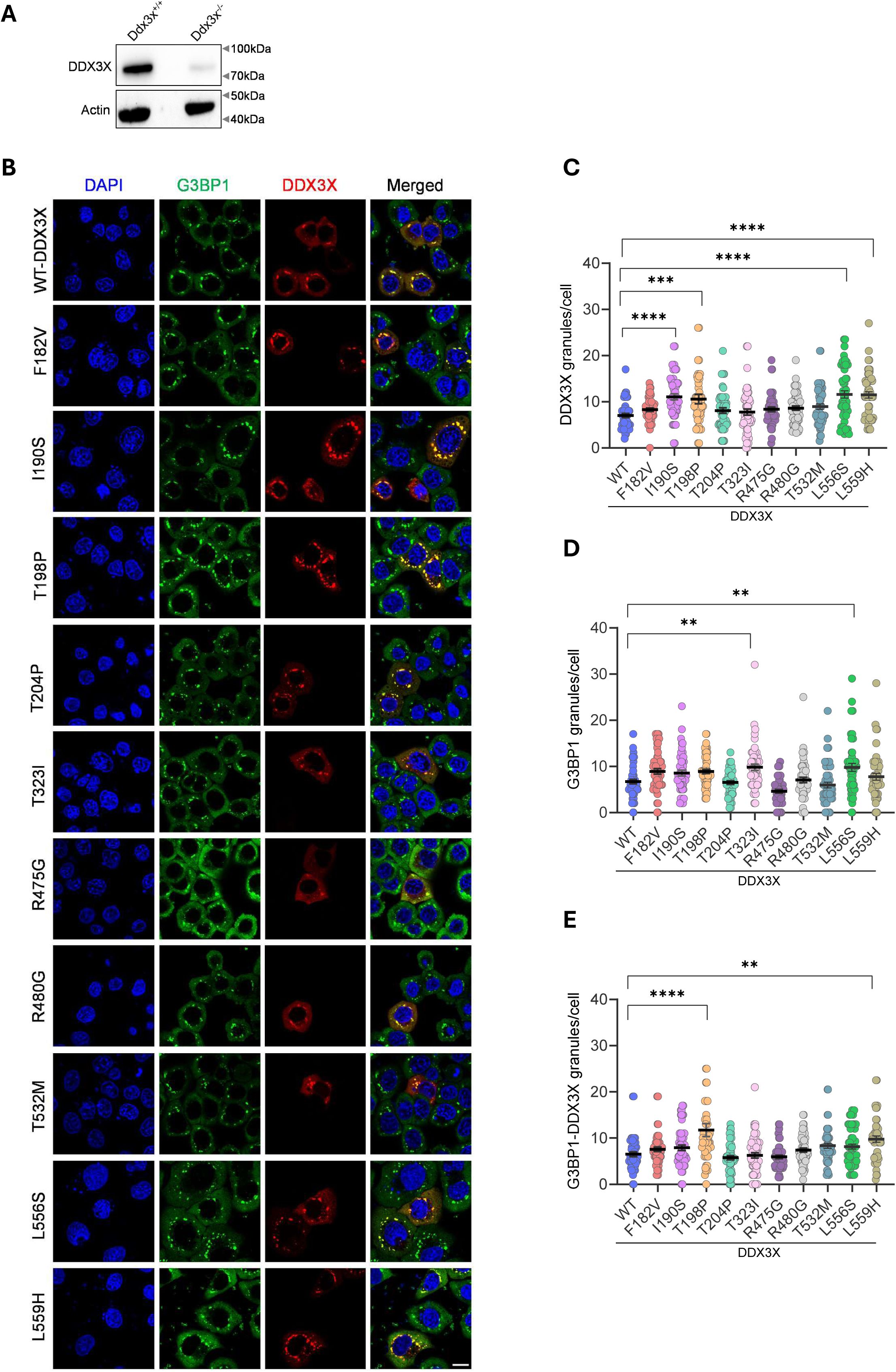
DDX3X missense mutations enhance stress granule assembly in HeLa cells. **A**. Immunoblot analysis showing levels of DDX3X protein in HeLa WT & Ddx3x^-/-^ cells. **B**. Representative confocal microscopy images of HeLa cells expressing WT or mutant DDX3X-mCherry constructs and subjected to SA (200µM) stress and immunostained for stress granule marker G3BP1 (green) and DAPI (blue). Scale bar=10µm. **C-E**. Quantification of numbers of DDX3X (red), G3BP1 (green), and DDX3X-G3BP1 (yellow) granules per cell quantified from ≥50 cells across three independent experiments exhibiting DDX3X colocalising with stress granules in HeLa cells. ****p < 0.0001, ***p = 0.0008 (DDX3X granules/cell), **p = 0.0022 (WT vs I190S), **p = 0.0028 (WT vs T323I), **p = 0.0031 (WT vs L556S) (G3BP1 granules/cells), ****p < 0.0001, **p = 0.0034 (G3BP1-DDX3X colocalised granules/cells) (One-way ANOVA test). Data shown are mean ± SEM.

**Figure S2.**
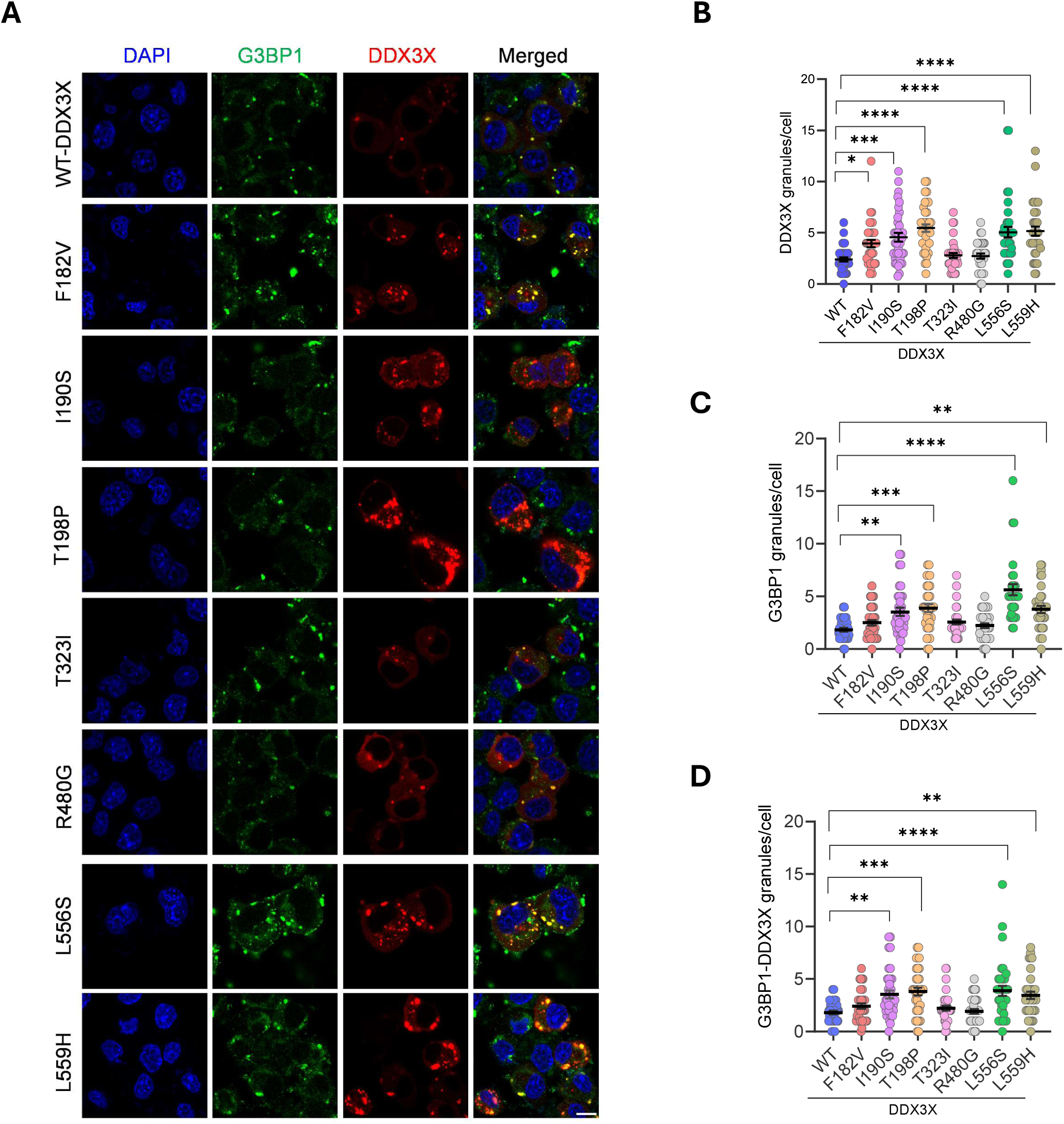
DDX3X missense mutations enhance assembly of stress granules in N2a cells. **A.** Representative confocal images of N2a cells expressing WT or mutant DDX3X-mCherry subjected to SA (100µM) stress for 2 hours and immunostained for stress granule marker G3BP1 (green) and DAPI (blue). Scale bar=10µm. **B-D**. Quantification of numbers of DDX3X (red), G3BP1 (green) and DDX3X-G3BP1 (yellow) granules per cell from ≥ 30cells across three independent experiments exhibiting DDX3X colocalising with stress granules in N2a cells. ****p < 0.0001, ***p = 0.0003, *p = 0.0181 (DDX3X granules/cell), ****p < 0.0001, ***p = 0.0001 (WT VS T198P), ***p = 0.0003 (WT vs L559H),**p = 0.0016 (G3BP1 granules/cells), ****p < 0.0001, ***p = 0.0001, **p = 0.0013 (WT vs I190S), **p = 0.0024 (WT vs L559H) (G3BP1-DDX3X colocalised granules/cells) (One-way ANOVA test). Data shown are mean ± SEM.

**Figure S3.**
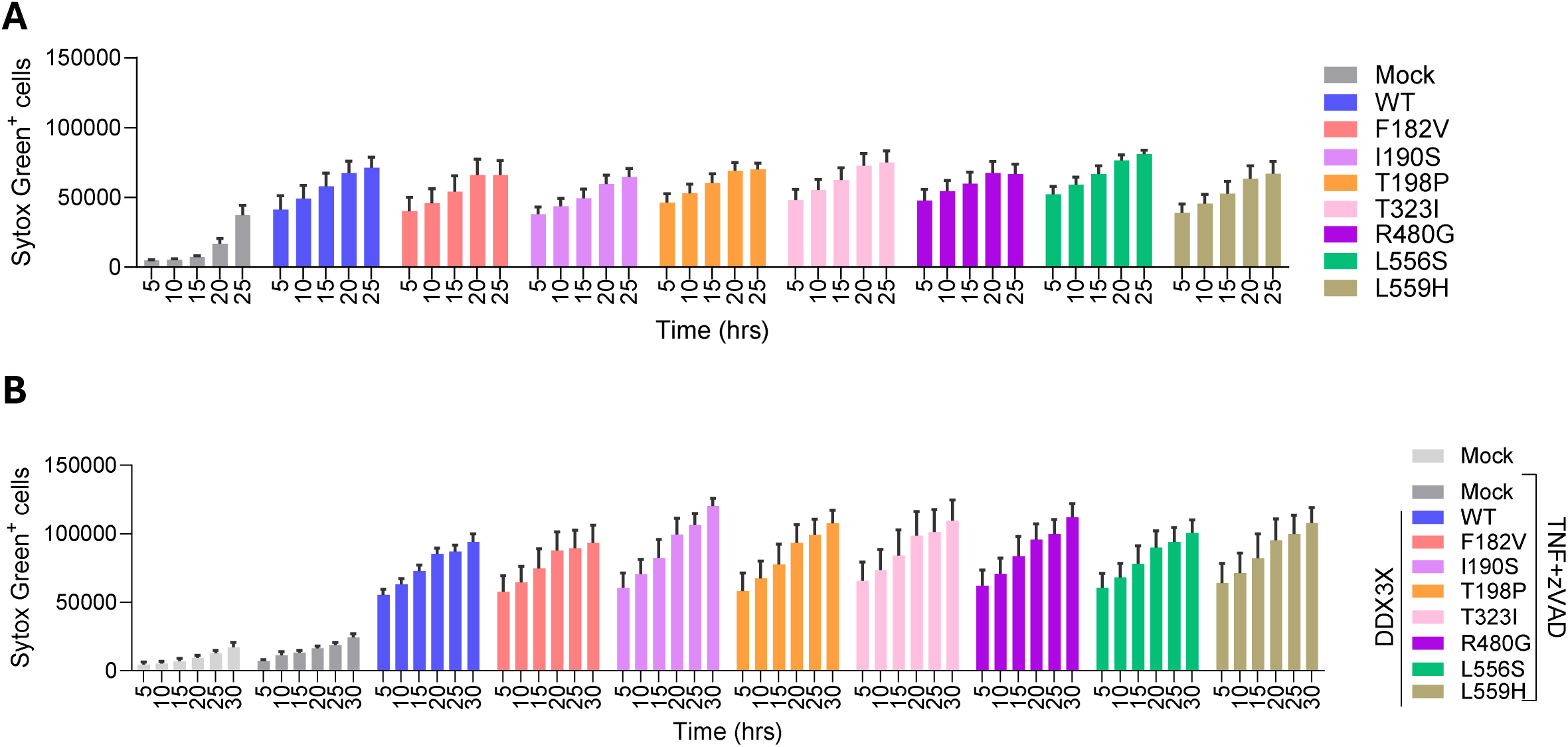
DDX3X missense mutations induced persistent granules enhance sensitivity to cytotoxic/cell death stimuli. **A.** Cell death measurement by Sytox green staining of N2a cells ectopically expressing WT or mutant DDX3X-mCherry. ns = non-significant (Two-way ANOVA test). Data shown are mean ± SEM. **B.** Cell death measurement by Sytox green staining of N2a cells ectopically expressing WT or mutant DDX3X-mCherry treated with TNF and zVAD. Ns = non-significant, (Two-way ANOVA test). Data shown are mean ± SEM.

**Figure S4.**
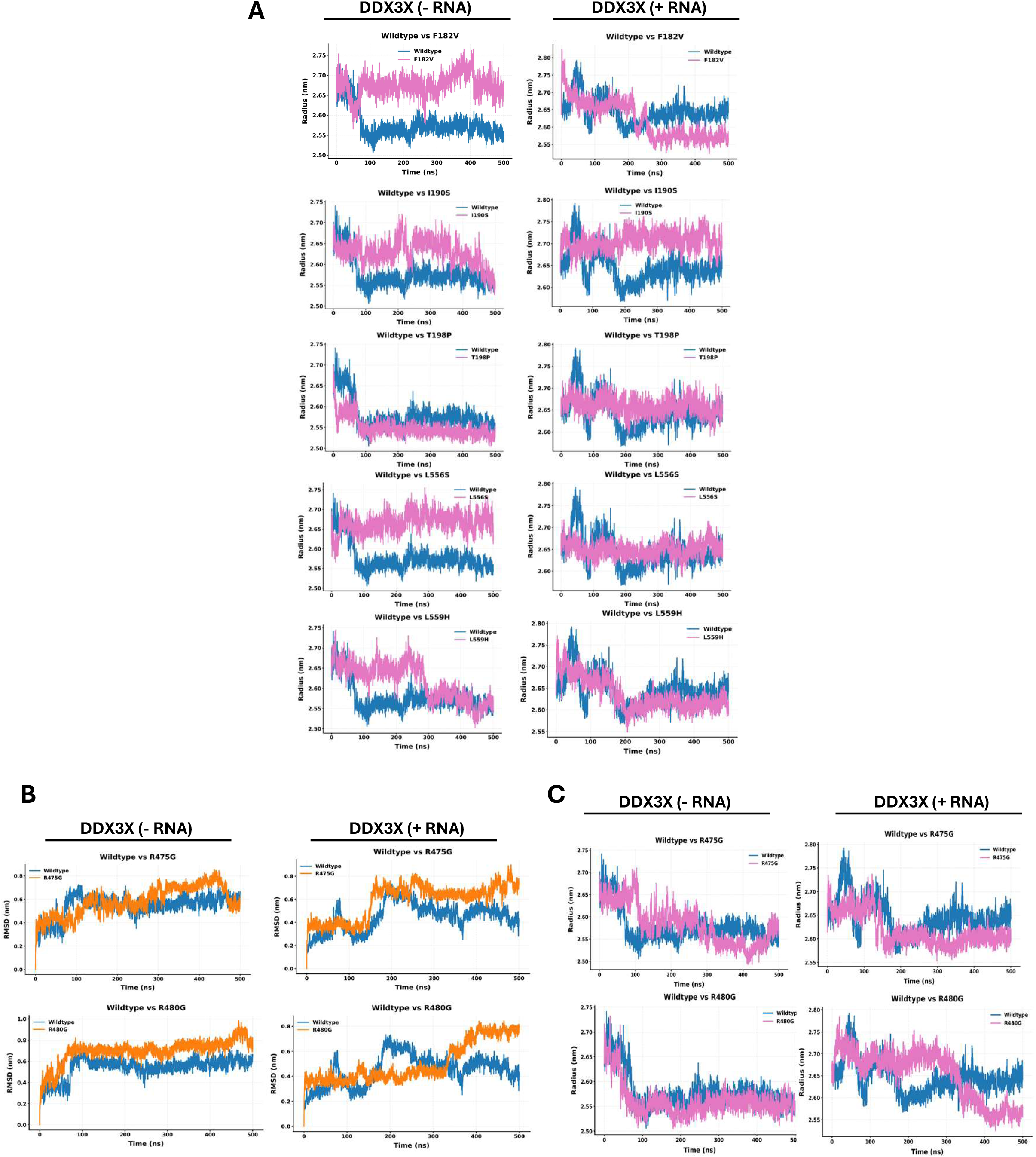
MD simulation showing Radius of Gyration and RMSD plots of selected DDX3X mutants. **A.** MD simulation-based Radius of gyration (Rg) plots showing the comparison of the WT-DDX3X trajectory (blue) against the indicated DDX3X mutant trajectories (magenta). **B-C.** MD simulations based RMSD (B) and Rg (C) trajectories of DDX3X mutants (R475G and R480G) (orange/magenta) in comparison with WT-DDX3X trajectory (blue).

**Fig. S5.**
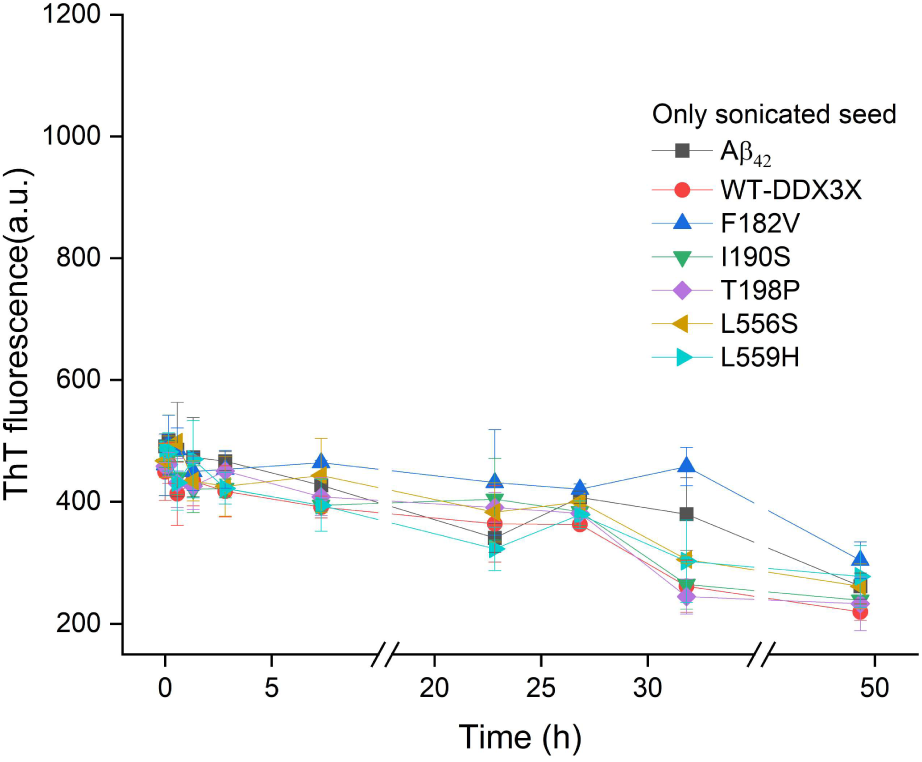
Time-dependent ThT fluorescence plots of sonicated DDX3X variant seeds without Aβ_1-42_ peptides, excited at 490nm.

## REFERENCES

1. Bol, G.M., Vesuna, F., Xie, M., Zeng, J., Aziz, K., Gandhi, N., Levine, A., Irving, A., Korz, D., Tantravedi, S., et al. (2015). Targeting DDX3 with a small molecule inhibitor for lung cancer therapy. EMBO Mol Med 7, 648–669.

2. Hilliker, A., Gao, Z., Jankowsky, E., and Parker, R. (2011). The DEAD-box protein Ded1 modulates translation by the formation and resolution of an eIF4F-mRNA complex. Mol Cell 43, 962–972.

3. Mo, J., Liang, H., Su, C., Li, P., Chen, J., and Zhang, B. (2021). DDX3X: structure, physiologic functions and cancer. Mol Cancer 20, 38.

4. Samir, P., Kesavardhana, S., Patmore, D.M., Gingras, S., Malireddi, R.K.S., Karki, R., Guy, C.S., Briard, B., Place, D.E., Bhattacharya, A., et al. (2019). DDX3X acts as a live-or-die checkpoint in stressed cells by regulating NLRP3 inflammasome. Nature 573, 590–594.

5. Shih, J.W., Wang, W.T., Tsai, T.Y., Kuo, C.Y., Li, H.K., and Wu Lee, Y.H. (2012). Critical roles of RNA helicase DDX3 and its interactions with eIF4E/PABP1 in stress granule assembly and stress response. Biochem J 441, 119–129.

6. Cruciat, C.M., Dolde, C., de Groot, R.E., Ohkawara, B., Reinhard, C., Korswagen, H.C., and Niehrs, C. (2013). RNA helicase DDX3 is a regulatory subunit of casein kinase 1 in Wnt-beta-catenin signaling. Science 339, 1436–1441.

7. Soulat, D., Burckstummer, T., Westermayer, S., Goncalves, A., Bauch, A., Stefanovic, A., Hantschel, O., Bennett, K.L., Decker, T., and Superti-Furga, G. (2008). The DEAD-box helicase DDX3X is a critical component of the TANK-binding kinase 1-dependent innate immune response. EMBO J 27, 2135–2146.

8. Kesavardhana, S., Samir, P., Zheng, M., Malireddi, R.K.S., Karki, R., Sharma, B.R., Place, D.E., Briard, B., Vogel, P., and Kanneganti, T.D. (2021). DDX3X coordinates host defense against influenza virus by activating the NLRP3 inflammasome and type I interferon response. J Biol Chem 296, 100579.

9. Samir, P., and Kanneganti, T.D. (2022). DEAD/H-Box Helicases in Immunity, Inflammation, Cell Differentiation, and Cell Death and Disease. Cells 11.

10. Valentin-Vega, Y.A., Wang, Y.D., Parker, M., Patmore, D.M., Kanagaraj, A., Moore, J., Rusch, M., Finkelstein, D., Ellison, D.W., Gilbertson, R.J., et al. (2016). Cancer-associated DDX3X mutations drive stress granule assembly and impair global translation. Sci Rep 6, 25996.

11. De Colibus, L., Stunnenberg, M., and Geijtenbeek, T.B.H. (2022). DDX3X structural analysis: Implications in the pharmacology and innate immunity. Curr Res Immunol 3, 100–109.

12. Song, H., and Ji, X. (2019). The mechanism of RNA duplex recognition and unwinding by DEAD-box helicase DDX3X. Nat Commun 10, 3085.

13. Molliex, A., Temirov, J., Lee, J., Coughlin, M., Kanagaraj, A.P., Kim, H.J., Mittag, T., and Taylor, J.P. (2015). Phase separation by low complexity domains promotes stress granule assembly and drives pathological fibrillization. Cell 163, 123–133.

14. Overwijn, D., and Hondele, M. (2023). DEAD-box ATPases as regulators of biomolecular condensates and membrane-less organelles. Trends Biochem Sci 48, 244–258.

15. Patmore, D.M., Jassim, A., Nathan, E., Gilbertson, R.J., Tahan, D., Hoffmann, N., Tong, Y., Smith, K.S., Kanneganti, T.D., Suzuki, H., et al. (2020). DDX3X Suppresses the Susceptibility of Hindbrain Lineages to Medulloblastoma. Dev Cell 54, 455–470.e455.

16. Gadek, M., Sherr, E.H., and Floor, S.N. (2023). The variant landscape and function of DDX3X in cancer and neurodevelopmental disorders. Trends Mol Med 29, 726–739.

17. Lennox, A.L., Hoye, M.L., Jiang, R., Johnson-Kerner, B.L., Suit, L.A., Venkataramanan, S., Sheehan, C.J., Alsina, F.C., Fregeau, B., Aldinger, K.A., et al. (2020). Pathogenic DDX3X Mutations Impair RNA Metabolism and Neurogenesis during Fetal Cortical Development. Neuron 106, 404–420.e408.

18. Kellaris, G., Khan, K., Baig, S.M., Tsai, I.C., Zamora, F.M., Ruggieri, P., Natowicz, M.R., and Katsanis, N. (2018). A hypomorphic inherited pathogenic variant in DDX3X causes male intellectual disability with additional neurodevelopmental and neurodegenerative features. Hum Genomics 12, 11.

19. Kennis, M.G.P., Rots, D., Bouman, A., Ockeloen, C.W., Boelen, C., Marcelis, C.L.M., de Vries, B.B.A., Elting, M.W., Waisfisz, Q., Suri, M., et al. (2025). DDX3X-related neurodevelopmental disorder in males - presenting a new cohort of 19 males and a literature review. Eur J Hum Genet 33, 980–988.

20. Kennis, M.G.P., Rots, D., Bouman, A., Ockeloen, C.W., Boelen, C., Marcelis, C.L.M., de Vries, B.B.A., Elting, M.W., Waisfisz, Q., Suri, M., et al. (2025). DDX3X-related neurodevelopmental disorder in males – presenting a new cohort of 19 males and a literature review. European Journal of Human Genetics 33, 980–988.

21. Snijders Blok, L., Madsen, E., Juusola, J., Gilissen, C., Baralle, D., Reijnders, M.R., Venselaar, H., Helsmoortel, C., Cho, M.T., Hoischen, A., et al. (2015). Mutations in DDX3X Are a Common Cause of Unexplained Intellectual Disability with Gender-Specific Effects on Wnt Signaling. Am J Hum Genet 97, 343–352.

22. Levy, T., Siper, P.M., Lerman, B., Halpern, D., Zweifach, J., Belani, P., Thurm, A., Kleefstra, T., Berry-Kravis, E., Buxbaum, J.D., et al. (2023). DDX3X Syndrome: Summary of Findings and Recommendations for Evaluation and Care. Pediatr Neurol 138, 87–94.

23. Radford, E.J., Tan, H.K., Andersson, M.H.L., Stephenson, J.D., Gardner, E.J., Ironfield, H., Waters, A.J., Gitterman, D., Lindsay, S., Abascal, F., et al. (2023). Saturation genome editing of DDX3X clarifies pathogenicity of germline and somatic variation. Nat Commun 14, 7702.

24. Hampel, H., Hardy, J., Blennow, K., Chen, C., Perry, G., Kim, S.H., Villemagne, V.L., Aisen, P., Vendruscolo, M., Iwatsubo, T., et al. (2021). The Amyloid-beta Pathway in Alzheimer’s Disease. Mol Psychiatry 26, 5481–5503.

25. Koo, E.H., Lansbury, P.T., Jr., and Kelly, J.W. (1999). Amyloid diseases: abnormal protein aggregation in neurodegeneration. Proc Natl Acad Sci U S A 96, 9989–9990.

26. Murray, D.T., Kato, M., Lin, Y., Thurber, K.R., Hung, I., McKnight, S.L., and Tycko, R. (2017). Structure of FUS Protein Fibrils and Its Relevance to Self-Assembly and Phase Separation of Low-Complexity Domains. Cell 171, 615–627 e616.

27. Villemagne, V.L., Burnham, S., Bourgeat, P., Brown, B., Ellis, K.A., Salvado, O., Szoeke, C., Macaulay, S.L., Martins, R., Maruff, P., et al. (2013). Amyloid beta deposition, neurodegeneration, and cognitive decline in sporadic Alzheimer’s disease: a prospective cohort study. Lancet Neurol 12, 357–367.

28. Cheng, J., Novati, G., Pan, J., Bycroft, C., Zemgulyte, A., Applebaum, T., Pritzel, A., Wong, L.H., Zielinski, M., Sargeant, T., et al. (2023). Accurate proteome-wide missense variant effect prediction with AlphaMissense. Science 381, eadg7492.

29. Owens, M.C., Shen, H., Yanas, A., Mendoza-Figueroa, M.S., Lavorando, E., Wei, X., Shweta, H., Tang, H.Y., Goldman, Y.E., and Liu, K.F. (2024). Specific catalytically impaired DDX3X mutants form sexually dimorphic hollow condensates. Nat Commun 15, 9553.

30. Guo, L., Kim, H.J., Wang, H., Monaghan, J., Freyermuth, F., Sung, J.C., O’Donovan, K., Fare, C.M., Diaz, Z., Singh, N., et al. (2018). Nuclear-Import Receptors Reverse Aberrant Phase Transitions of RNA-Binding Proteins with Prion-like Domains. Cell 173, 677–692 e620.

31. Mateju, D., Franzmann, T.M., Patel, A., Kopach, A., Boczek, E.E., Maharana, S., Lee, H.O., Carra, S., Hyman, A.A., and Alberti, S. (2017). An aberrant phase transition of stress granules triggered by misfolded protein and prevented by chaperone function. EMBO J 36, 1669–1687.

32. Wolozin, B., and Ivanov, P. (2019). Stress granules and neurodegeneration. Nat Rev Neurosci 20, 649–666.

33. Calviello, L., Venkataramanan, S., Rogowski, K.J., Wyler, E., Wilkins, K., Tejura, M., Thai, B., Krol, J., Filipowicz, W., Landthaler, M., et al. (2021). DDX3 depletion represses translation of mRNAs with complex 5’ UTRs. Nucleic Acids Res 49, 5336–5350.

34. Ryan, C.S., and Schroder, M. (2022). The human DEAD-box helicase DDX3X as a regulator of mRNA translation. Front Cell Dev Biol 10, 1033684.

35. Reineke, L.C., Cheema, S.A., Dubrulle, J., and Neilson, J.R. (2018). Chronic starvation induces noncanonical pro-death stress granules. J Cell Sci 131.

36. Zhang, P., Fan, B., Yang, P., Temirov, J., Messing, J., Kim, H.J., and Taylor, J.P. (2019). Chronic optogenetic induction of stress granules is cytotoxic and reveals the evolution of ALS-FTD pathology. Elife 8.

37. Siddu, A., Natale, S., Wong, C.H., Shaye, H., and Südhof, T.C. (2025). Aggregation shifts amyloid-β peptides from synaptogenic to synaptotoxic. J Clin Invest 135.

38. Bhardwaj, T., Gadhave, K., Kapuganti, S.K., Kumar, P., Brotzakis, Z.F., Saumya, K.U., Nayak, N., Kumar, A., Joshi, R., Mukherjee, B., et al. (2023). Amyloidogenic proteins in the SARS-CoV and SARS-CoV-2 proteomes. Nat Commun 14, 945.

39. De Felice, F.G., Velasco, P.T., Lambert, M.P., Viola, K., Fernandez, S.J., Ferreira, S.T., and Klein, W.L. (2007). Abeta oligomers induce neuronal oxidative stress through an N-methyl-D-aspartate receptor-dependent mechanism that is blocked by the Alzheimer drug memantine. J Biol Chem 282, 11590–11601.

40. Kadowaki, H., Nishitoh, H., Urano, F., Sadamitsu, C., Matsuzawa, A., Takeda, K., Masutani, H., Yodoi, J., Urano, Y., Nagano, T., et al. (2005). Amyloid beta induces neuronal cell death through ROS-mediated ASK1 activation. Cell Death Differ 12, 19–24.

41. Luciunaite, A., McManus, R.M., Jankunec, M., Racz, I., Dansokho, C., Dalgediene, I., Schwartz, S., Brosseron, F., and Heneka, M.T. (2020). Soluble Abeta oligomers and protofibrils induce NLRP3 inflammasome activation in microglia. J Neurochem 155, 650–661.

42. Salvadores, N., Moreno-Gonzalez, I., Gamez, N., Quiroz, G., Vegas-Gomez, L., Escandon, M., Jimenez, S., Vitorica, J., Gutierrez, A., Soto, C., et al. (2022). Abeta oligomers trigger necroptosis-mediated neurodegeneration via microglia activation in Alzheimer’s disease. Acta Neuropathol Commun 10, 31.

43. Hondele, M., Sachdev, R., Heinrich, S., Wang, J., Vallotton, P., Fontoura, B.M.A., and Weis, K. (2019). DEAD-box ATPases are global regulators of phase-separated organelles. Nature 573, 144–148.

44. Yanas, A., Shweta, H., Owens, M.C., Liu, K.F., and Goldman, Y.E. (2024). RNA helicases DDX3X and DDX3Y form nanometer-scale RNA-protein clusters that support catalytic activity. Curr Biol 34, 5714–5727.e5716.

45. Li, Q., Zhang, P., Zhang, C., Wang, Y., Wan, R., Yang, Y., Guo, X., Huo, R., Lin, M., Zhou, Z., et al. (2014). DDX3X regulates cell survival and cell cycle during mouse early embryonic development. J Biomed Res 28, 282–291.

46. Mosti, F., Hoye, M.L., Escobar-Tomlienovich, C.F., and Silver, D.L. (2025). Multi-modal investigation reveals pathogenic features of diverse DDX3X missense mutations. PLoS Genet 21, e1011555.

47. de Castro Fonseca, M., de Oliveira, J.F., Araujo, B.H.S., Canateli, C., do Prado, P.F.V., Amorim Neto, D.P., Bosque, B.P., Rodrigues, P.V., de Godoy, J.V.P., Tostes, K., et al. (2021). Molecular and cellular basis of hyperassembly and protein aggregation driven by a rare pathogenic mutation in DDX3X. iScience 24, 102841.

48. Rio, D.C., Ares, M., Jr., Hannon, G.J., and Nilsen, T.W. (2010). Purification of RNA using TRIzol (TRI reagent). Cold Spring Harb Protoc 2010, pdb.prot5439.

